# A widespread family of ribosomal peptide metallophores involved in bacterial adaptation to metal stress

**DOI:** 10.1101/2024.03.18.585515

**Authors:** Laura Leprevost, Sophie Jünger, Guy Lippens, Céline Guillaume, Giuseppe Sicoli, Lydie Oliveira, Alex Rivera-Millot, Gabriel Billon, Céline Henry, Rudy Antoine, Séverine Zirah, Svetlana Dubiley, Yanyan Li, Françoise Jacob-Dubuisson

**Author notes:** contributed equally (Laura Leprevost, Sophie Jünger). INRS-Centre Armand-Frappier Santé Biotechnologie, Bacterial Symbionts Evolution, Laval, Quebec H7V 1B7, Canada. Author contributions: GL, RA, SD, SZ, YL, FJD designed research; LL, SJ, GL, CG, GS, LO, ARM, GB, CH, RA, SD performed research; LL, SJ, GL, CG, GS, LO, ARM, GB, CH, RA, SZ, SD, YL, FJD analyzed data; LL, SJ, GL, SD, SZ, YL, FJD wrote the paper. **Competing interest statement**: The authors declare no competing interests. **Classification:** Biological Sciences - Biochemistry - Microbiology.

## Abstract

Ribosomally synthesized and post-translationally modified peptides (RiPPs) are a structurally diverse group of natural products that bacteria employ in their survival strategies. Herein, we characterized the structure, the biosynthetic pathway and the mode of action of a new RiPP family called bufferins. With thousands of homologous biosynthetic gene clusters throughout the eubacterial phylogenetic tree, bufferins form by far the largest family of RiPPs modified by multinuclear non-heme iron-dependent oxidases (MNIO, DUF692 family). Using *Caulobacter vibrioides* bufferins as a model, we showed that the conserved Cys residues of their precursors are transformed into 5-thiooxazoles, further expanding the reaction range of MNIO enzymes. This rare modification is installed in conjunction with a partner protein of the DUF2063 family. Bufferin precursors are the first examples of bacterial RiPPs found to feature an N-terminal Sec signal peptide and thus to be exported by the ubiquitous Sec pathway, a new paradigm in the RiPP field. Other original features of bufferins are their large size and protein-like fold, which blurs the line between modified peptides and proteins. We reveal that bufferins are involved in copper homeostasis, and their metal-binding propensity requires the thiooxazole heterocycles. Bufferins enhance bacterial growth under copper stress by sequestering excess metal ions in the periplasm. Our study thus describes a large family of RiPP metallophores and unveils a widespread but overlooked metal homeostasis mechanism in eubacteria likely to be relevant to One-Health strategies.

**Significance statement:** Copper is both essential and toxic in excess. Bacteria face copper in their environments, notably in phagocytes, hence they have developed several defense mechanisms. We discovered a widespread strategy of protection from copper, through the biosynthesis of natural products that we call bufferins. Bufferins are <u>ri</u>bosomally synthesized <u>p</u>ost-translationally modified <u>p</u>eptides (RiPPs), natural products with key roles in bacterial physiology and ecology. Bufferins enhance bacterial growth under copper stress by complexing with the metal using thiooxazole heterocycles that result from enzymatic modification of cysteine residues. With thousands of homologs throughout the eubacterial phylogenetic tree, bufferins represent a highly prevalent strategy of adaptation to metal stress. They are larger in size than most RiPPs, expanding the concept of RiPPs to modified proteins.

## Introduction

Ribosomally synthesized and post-translationally modified peptides (RiPPs) are natural products with a tremendous variety of structures and biological activities (1–3). They are synthesized from a gene-encoded precursor peptide, generally composed of an N-terminal leader sequence and a core sequence where post-translational modifications (PTMs) occur. In many cases, a pathway-specific RiPP recognition element (RRE) domain interacts with the leader sequence and mediates recruitment of the modification enzymes (4–6). RRE domains are thus a hallmark of RiPP biosynthesis. The vast structural diversity of RiPPs is conferred by virtually unlimited possibilities of chemical transformations. An emerging family of PTM enzymes is the DUF692 family of multinuclear non-heme iron dependent oxidases (MNIOs) that catalyze various reactions in the biosynthesis of diverse RiPPs, mostly but not exclusively on Cys residues (7–13). These reactions include oxazolone coupled to thioamide formation, excision of cysteine β-methylene, Cys-involved macrocyclization, and C-C and C-N cleavages (7, 8, 11–13). However, the full spectrum of MNIO activities remains far from being fully appreciated.

Four-gene *gig* operons (*gig* for *<u>g</u>*old-*i*nduced *<u>g</u>*enes) encoding DUF692-family proteins were identified in the genomes of several microbial species (*SI* Fig. S1). In *Cupriavidus metallidurans*, *Legionella pneumophila*, and *Caulobacter vibrioides*, the expression of these operons is induced under various transition metal stresses that include gold, cadmium, copper, etc., whereas in *Bordetella pertussis* it is controlled by the master regulator of virulence BvgAS (14–18). Orthologous operons were notably found in methane-oxidizing bacteria (19). However, despite genetic evidence for an involvement in the copper-resistant phenotype in *C. metallidurans* (20), the function of these operons remains elusive. Based on their composition, we speculated that they are biosynthetic gene clusters (BGCs) whose products could represent a new RiPP family. Unlike all previously described RiPPs however, these putative precursors have Sec signal peptides. They are also markedly larger than known MNIO-modified RiPP precursors. Using genetic, structural, and bioinformatic analyses, we questioned the function, the mode of action, and the prevalence of *gig*-like operons in bacteria.

We report the characterization of the products of the *C. vibrioide*s *gig*-like BGCs, the first representative of bufferins, a new family of RiPPs. We revealed that bufferins are small protein metallophores containing a rare 5-thiooxazole modification of cysteine residues. The joint action of the DUF692 (MNIO) and DUF2063 proteins is required to install this post-translational modification, which is new to the MNIO family of enzymes. We showed that the *C. vibrioides* bufferins protect bacteria against toxic concentrations of copper by sequestering the excess metal in the periplasm. Our bioinformatic study suggests that bufferins represent an overlooked transition metal defense mechanism very common in Hydrobacteria and Terrabacteria.

## Results

### Large new RiPP family involving MNIOs

We performed an *in-silico* analysis of the CCNA_03363-03366 operon of *C. vibrioides*, hereafter named *buf1*, to gain insight into gene functions. CCNA_03363 encodes a DUF2282-family protein (BufA1), whose members are 80-120 residues long and harbor four Cys residues (Cys^I^ to Cys^IV^) and a predicted Sec signal peptide (Fig. 1A). We hypothesized that BufA1 is an unusually long precursor peptide whose translocation across the cytoplasmic membrane would rely on the ubiquitous Sec export pathway rather than on a specialized export system as is often the case for secreted RiPPs (21). The AlphaFold2 model predicts that the 68-residue long BufA1 core peptide contains a 2-stranded β sheet and some α-helical structure (*SI* Fig. S1). It thus folds into a small protein rather than being a totally unstructured peptide.

**Figure 1.**
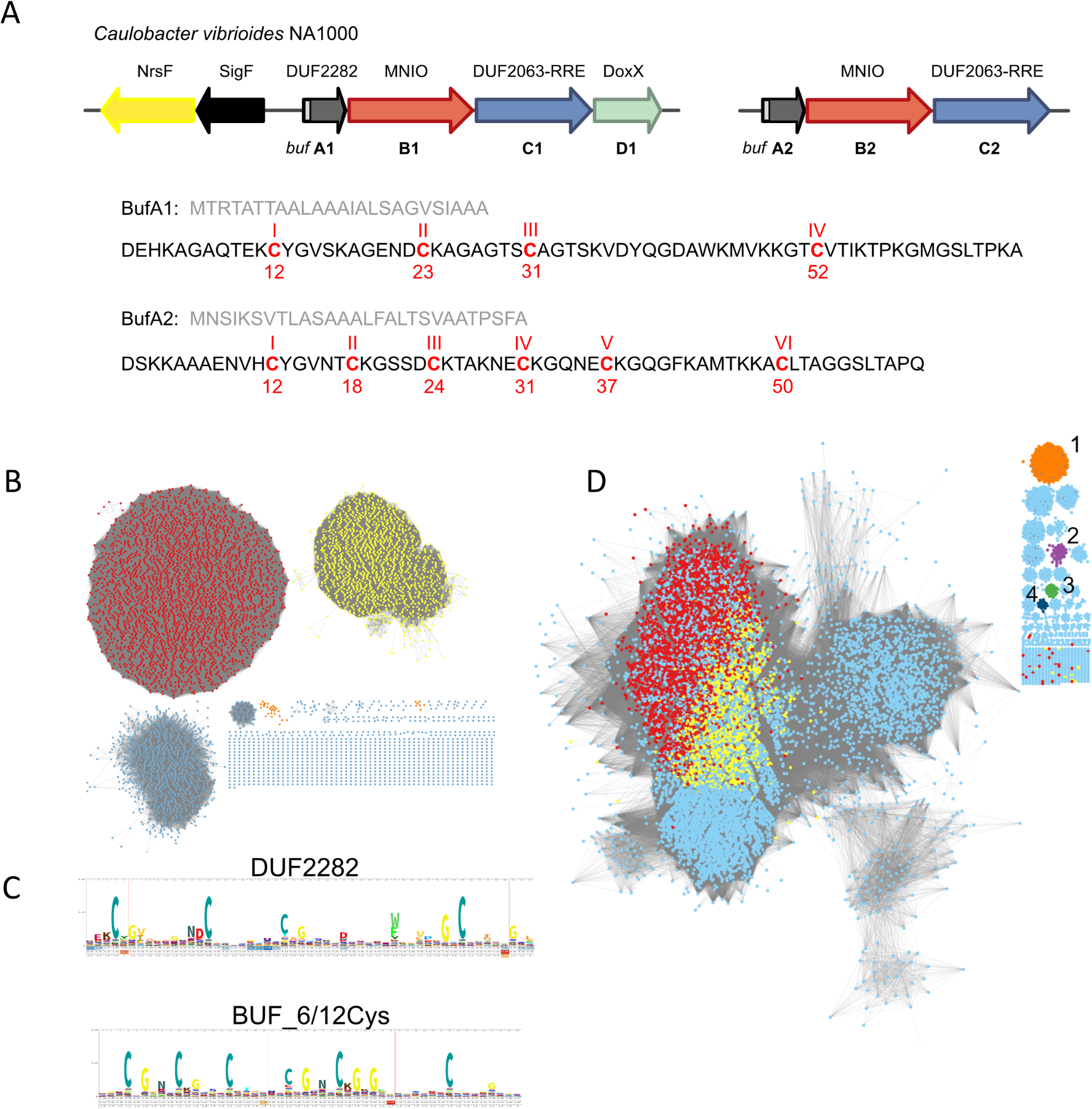
*In silico* analyses of the bufferin family. **A**. Gene composition of the *buf1* and *buf2* BGCs and sequences of BufA1 and BufA2 with the Cys residues labeled (numbering according to the core peptides). The signal peptide and core peptide sequences are in pale and dark grey, respectively. In the vicinity of *buf1ABCD* are genes coding for SigF and the anti-sigma factor NrsF that regulate *buf1* and *buf2* expression. **B**. SSN analysis of bufferin-like precursors genetically associated with MNIOs. DUF2282 and BUF_6/12Cys proteins, including BufA1 and BufA2 of *C. vibrioides*, belong to the largest two clusters (red and yellow dots), respectively. Small clusters in orange contain the RiPPs of *Chryseobacterium* (7) that are predicted to harbor signal peptides. **C**. Weblogos for the DUF2282 and BUF_6/12Cys bufferins. **D**. SSN analysis of MNIOs (at 80% identity). MNIOs genetically associated with DUF2282 and BUF_6/12Cys proteins, including BufB1 and BufB2 of *C. vibrioides*, are in red and yellow, respectively. MNIOs associated with PTMS on methanobactins, *Chryseobacterium* RiPPs, TglA-type pearlins and aminopyruvatides are found in small clusters numbered 1 to 4.

The product of CCNA_03364 (BufB1) belongs to the MNIO family, whereas CCNA_03365 codes for a protein (BufC1) with an uncharacterized N-terminal DUF2063 domain. The predicted structural model of BufC1 revealed that its C-terminal domain is related to RRE domains (4–6) (*SI* Fig. S1). In light of recent discoveries of MNIOs involved in RiPP biosynthesis (7–13), we hypothesized that BufB1 and BufC1 would install modifications on Cys residues of BufA1. Finally, CCNA_03366 encodes a putative DoxX-type oxido-reductase, BufD1 (22, 23).

Inspection of the *C. vibrioides* genome identified a second operon, the *buf2* BGC (CCNA_02999-03001) (Fig. 1A) which encodes homologs of BufB1 (CCNA_03000) and BufC1 (CCNA_02999) but lacks a *bufD1*-like gene. Like BufA1, its putative precursor peptide BufA2 (CCNA_03001) possesses a Sec signal peptide. Its 61-residue core peptide contains six Cys residues (Cys^I^ to Cys^VI^) (Fig. 1A). Although its predicted 3D structure is very similar to that of BufA1, this small protein does not have a corresponding profile in the databases of protein families (*SI* Fig. S1).

To evaluate the abundance of homologous BGCs, we collected >14,000 MNIO protein sequences from the NCBI non-redundant database. The retrieved sequences are widely distributed throughout Pseudomonadota, Terrabacteria, Myxococcota, Fibrobacteres/Chlorobi/Bacterioidetes, Planctomycetes/Verrucomicrobia/Chlamydia, and Acidobacteriota. We analyzed their genomic contexts and collected associated genes coding for putative bufferin-like precursors, i.e., small proteins with Cys residues and a predicted Sec signal peptide. Thousands of these were collected, some of which are markedly larger than typical RiPP precursors and harbor numerous Cys residues (detailed *in silico* analyses will be published separately). In a sequence similarity network (SSN) constructed with these proteins, the largest cluster corresponds to the DUF2282 signature, including *C. vibrioides* BufA1 (Fig. 1B). BufA2 of *C. vibrioides* belongs to the second largest cluster of proteins characterized by six or twelve conserved Cys residues, which we named BUF_6/12Cys (Fig. 1B,C). Of note, BUF_6/12Cys proteins were previously identified in Antarctic soil metagenomes (24). Parallel SSN analyses of the MNIO superfamily (Fig. 1D) showed that those coded in the DUF2282 and BUF_6/12Cys BGCs belong to the most populated sequence cluster, distinct from small clusters of MNIOs involved in the biosynthesis of other RiPP types (7, 9–11, 13). Together our data indicate that bufferins currently form the largest group of MNIO-associated RiPPs.

### Bufferins mediate adaptation to copper stress

In *C. vibrioides*, *buf1* and *buf2* BGCs belong to the SigF-controlled, core regulon of copper stress (15, 17, 25). To test if they could be up-regulated by other biologically relevant metals, we generated reporter fusions within *bufA1* and *bufA2* and confirmed that both are induced by high concentrations of Cu^2+^, but not by Fe^2+^, Zn^2+^ or Mn^2+^ (Fig. 2A; *SI* Fig. S2).

**Figure 2.**
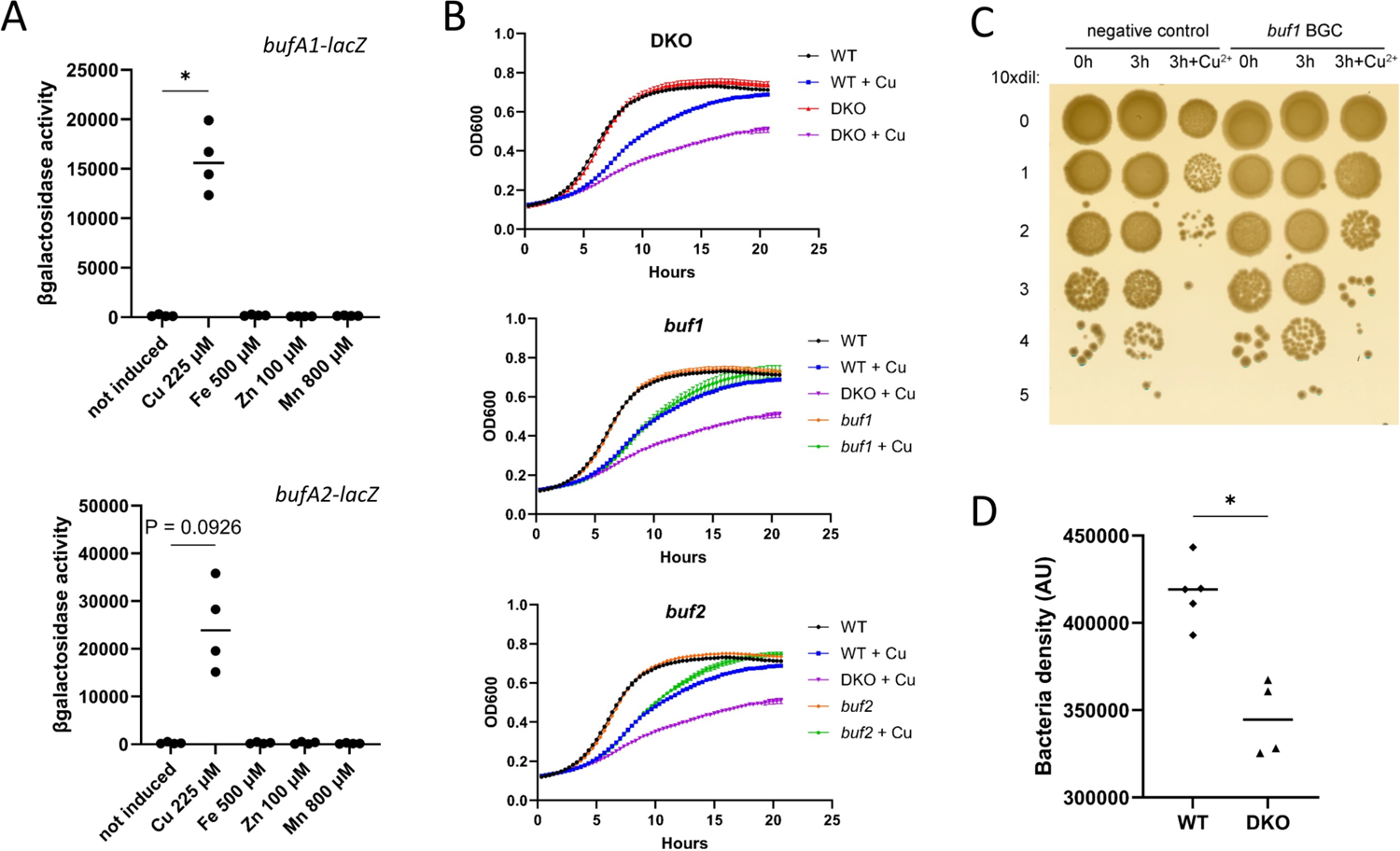
Regulation and function of the bufferins. **A**. Reporter assays with *bufA1-lacZ* and *bufA2-lacZ* transcriptional fusions. Bacteria were grown for 16 h with the indicated concentrations of CuSO_4_, FeSO_4_, ZnSO_4_ or MnCl_2_. Non-parametric, one-way analysis of variance Kruskal-Wallis (two-sided) tests followed by a Dunn’s multiple-comparison tests were used to analyze the differences between treated cultures and the control (n=4; * in the upper left graph indicates p=0.047). **B**. Effect of copper on bacterial growth. The Cv_DKO_, Cv*_buf1ABCD_* (*buf1)* and Cv*_buf2ABC_* (*buf2*) strains were grown without or with 225 µM CuSO_4_. **C**. Gain-of-function assay. Expression of *buf1* in *E. coli* BL21(pCA24-buf1A^str^BCD) was induced by IPTG, and the bacteria were challenged with 0.3 mM CuSO_4_ for three hours before serial dilutions. BL21(pCA24-psmCA) expressing a non-relevant RiPP BGC was used as a control. **D**. Assay of bacterial lysis by *D. discoidum*. Bacterial densities of the lysis plaques were determined using ImageJ analyses. A non-parametric Mann-Whitney test (two-sided) was used to analyze the differences between the Cv_WT_ strain (n=5) and the Cv_DKO_ strain (n=4; * indicates p = 0.0159). The medians are shown.

To determine if bufferins, the putative RiPP products of the *buf* BGCs, contribute to adaptation to copper stress, we prepared *C. vibrioides* deletion mutants that express a single (Cv_buf1_ and Cv_buf2_) or neither BGC (Cv_DKO_). Cv_DKO_ grew more slowly than the parental Cv_WT_ strain when CuSO_4_ was added to the growth medium, unlike Cv_buf1_ or Cv_buf2_ (Fig. 2B). Thus, bufferins enhance growth under copper stress and are functionally redundant. To confirm that a single BGC is sufficient to protect bacteria from excess copper, we performed a gain-of-function experiment. Expression of *buf1ABCD* in an *Escherichia coli* laboratory strain enhanced survival of bacteria exposed to a bactericidal concentration of CuSO_4_ (Fig. 2C).

We then asked whether the bufferins would confer an advantage on *C. vibrioides* in biotic interactions. Since amoeba notably use copper against their bacterial preys (26), a predation assay was performed. We attempted to measure survival of *C. vibrioides* after ingestion by *Dictyostelium discoidum*, a well-established mode organism for the study of phagocytosis. However, as these assays were unsuccessful, we tested if the presence of the *buf* BGCs might slow down bacterial killing. We placed amoebal suspensions onto bacterial lawns, and after incubation we determined bacterial densities in the contact zones by densitometry scanning. Larger zones of bacterial clearance were observed for the Cc_DKO_ than for the Cc_WT_ strain, consistent with a protective role of bufferins in this context (Fig. 2D).

### Characterization of bufferin PTMs

To identify the PTMs of bufferins, we engineered an IPTG-inducible version of SigF to control expression of their BGCs in *C. vibrioides*. The metabolic profiles of IPTG-treated Cv_buf1_(pSigF) and Cv_buf2_(pSigF) were analyzed by liquid chromatography coupled to mass spectrometry (LC-MS) and compared to that of Cv_DKO_(pSigF) to reveal peptide species only present in the *buf*-expressing strains. In Cv_buf1_(pSigF), a major species exhibited a mass matching that of BufA1 with the Sec signal peptide cleaved off and a −10 Da mass shift (Fig. 3A; *SI* Fig. S3 and Table S1), suggesting that the core peptide accumulates in the periplasm and contains post-translational modifications of one or more residues. A second form with a −12 Da mass shift was also detected, but not systematically, and it disappeared upon freeze-thawing of cell extracts. The most stable product with a −10 Da mass shift is considered the mature peptide product and was named bufferin 1 (Buf1). Similarly, Cv_buf2_(pSigF) produced a major peptide product with a −18 Da mass shift, called bufferin 2 (Buf2), and a minor species with a −20 Da mass shift (*SI* Fig. S3 and Table S1).

**Figure 3.**
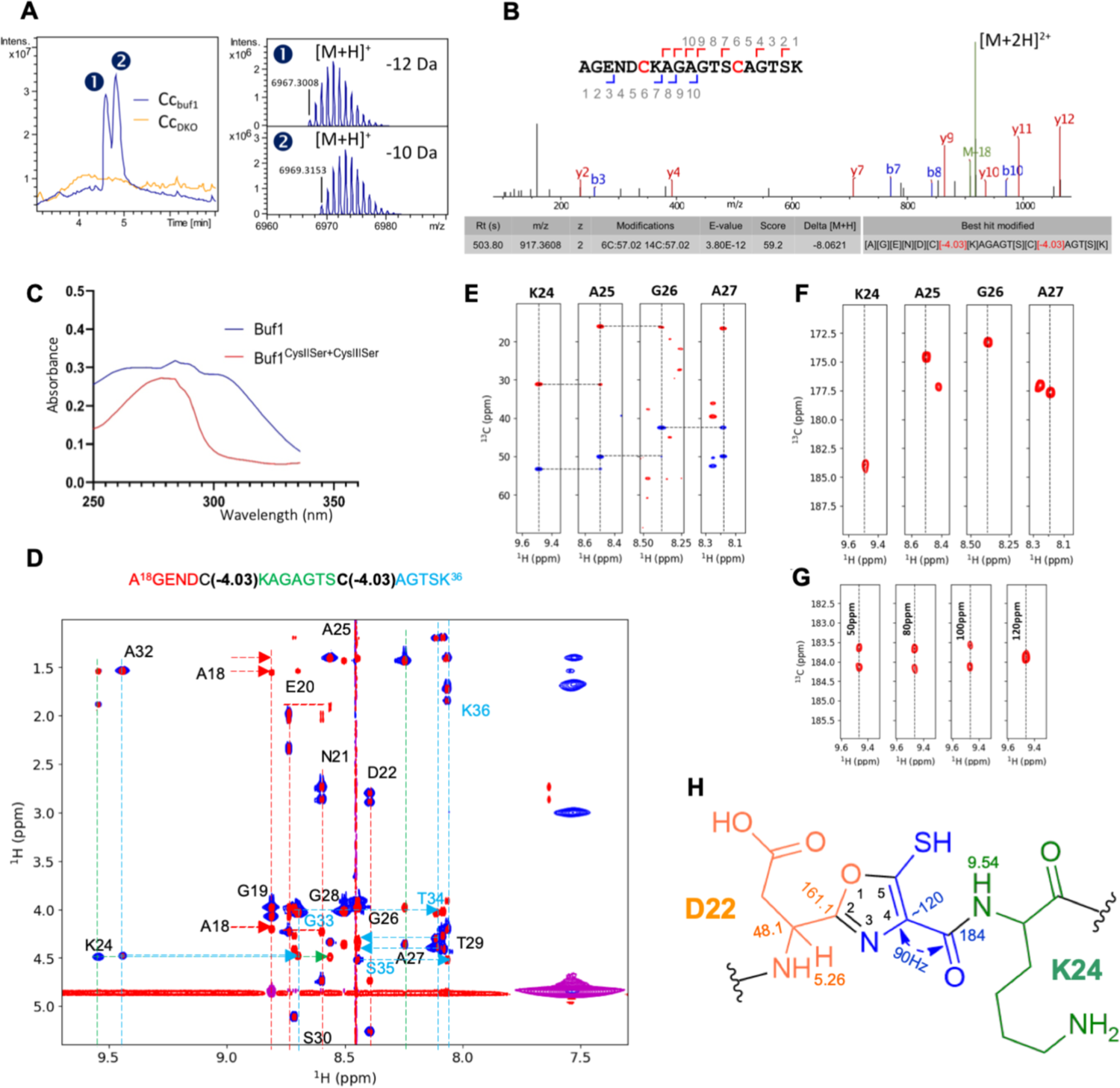
Post-translational modifications of bufferin 1. **A.** Top-down LC-MS analyses of cell extracts of Cv_buf1_(pSigF) compared with Cv_DKO_(pSigF). Left panel: total ion chromatograms, right panels: deconvoluted mass spectra of the compounds detected for Cv_buf1_(pSigF). Their Mw correspond to the core peptide (*i.e.*, without signal peptide; calculated monoisotopic Mw=6978.35 Da), with mass shifts of −10 Da ([M+H]^+^ at *m/z* 6969.32) and −12 Da ([M+H]^+^ at *m/*z 6967.30). **B.** Bottom-up analysis of Buf1. MS/MS spectrum of the central tryptic peptide ([M+2H]^2+^ at *m/z* 917.36). The two Cys residues carry carbamidomethyl groups resulting from alkylation with iodoacetamide (+ 57.02 Da), together with −4.03 Da mass shifts. **C.** UV/vis spectra of affinity-purified Buf1 and the Cys^II^Ser+Cys^III^Ser mutant. **D.** NMR ROESY/TOCSY experiments to assign the resonances of the 19-mer peptide. **E**. HNCACB experiment on the uniformly ^13^C, ^15^N-labeled peptide: sequential walk for the sequence following Cys^II^. **F**. The HNCO planes at the ^15^N frequency of the indicated residues show classical ∼175-ppm values for all carbonyl resonances, except for that downstream of a modified Cys, whose carbonyl carbon resonates at 184 ppm. **G**. HNCO planes through the resonance of Lys24 that follows Cys^II^ while modifying the offset of the ^13^C decoupling pulse during the C=O evolution period. Only when centered at 120 ppm is the 90-Hz carbon-carbon coupling refocused. **H.** Measured NMR parameters and proposed structure for the Buf1 post-translational modifications (shown for Cys^II^).

To test the hypothesis that mature bufferins contain modified Cys residues and disulfide (S-S) bonds, we performed reduction and alkylation treatment of the peptides. This yielded species carrying three or four alkylated groups for Buf1 and four, five or six alkylated groups for Buf2 (*SI* Fig. S4), suggesting that the expected PTMs do not affect Cys residues or result in alkylation-sensitive groups. MS analyses of trypsin-digested, alkylated bufferins revealed that Cys^I^ (Cys12) and Cys^IV^ (Cys52) in Buf1 (numbering according to the core peptide, see Fig. 1) had mass increments corresponding to alkylation only, indicative of unmodified residues (*SI* Fig. S5). By contrast, in the central tryptic peptide containing Cys^II^ (Cys23) and Cys^III^ (Cys31), both Cys residues carry mass increments corresponding to alkylation with additional −4.03 Da mass shifts (Fig. 3B), as proposed by the untargeted PTM search solution SpecGlobX (27).

To confirm that Cys residues are required for Buf1 function, we constructed strains with the conserved Cys residues replaced with Ser, separately or in combinations. Consistent with the proposed modifications of Cys^II^ or Cys^III^, the substitution of either of them yielded peptides with mass shifts of −6 Da relative to the calculated mass of the mutated peptide (*SI* Fig. S6). We tested the ability of the mutated BufA1 to protect *C. vibrioides* from excess copper. All mutant strains grew more slowly than the parental Cv_buf1_ in the presence of CuSO_4_ (*SI* Fig. S7), indicating the importance of modified and unmodified Cys residues for function. Altogether, the data are consistent with the presence of an S-S bond between Cys^I^ and Cys^IV^ and Cys^II^ and Cys^III^ being targets of 4-electron oxidation reactions that generate alkylation-sensitive groups. The Buf1 congener product with a −12 Da mass shift would correspond to a form with a labile S-S bond between the modified residues. Similarly, in Buf2, in addition to the unmodified Cys^I^ (Cys24) and Cys^VI^ (Cys50) likely forming an S-S bond (*SI* Fig. S1), the central Cys residues carry −4.03 Da shifts (*SI* Fig. S8).

To facilitate bufferin purification and characterization, we added C-terminal tags to BufA1 (*SI* Table S1). The affinity-purified tagged Buf1 was fully modified (*SI* Fig. S9). To probe if the modified Cys residues may be part of heterocycles as in other MNIO-modified RiPPs (7, 10), we recorded UV-visible absorption spectra of Buf1 and the Buf1^CysIISer+CysIIISer^ variant. Consistent with the presence of aromatic heterocycles (28), the spectrum of the former showed a broad absorbance peak extending to 310 nm, unlike that of the latter (Fig. 3C). The spectrum of recombinant Buf2 showed a similar absorbance maximum around 305 nm (*SI* Fig. S10).

### PTM identification by NMR

To increase the yield of bufferin for structural elucidation by NMR, we co-expressed *bufA1* encoding an N-terminal His_6_-SUMO-tag with *bufB1* and *bufC1* in *E. coli*. Purification of modified BufA1 followed by trypsin digestion and preparative HPLC allowed the preparation of the Cys^II^- and Cys^III^-containing 19-mer peptide at a milligram scale (*SI* Fig. S11). Its ^1^H spectrum indicated two groups of anomalous resonances (*SI* Fig. S12). By combining homonuclear ROESY and TOCSY experiments (29), they were assigned to the amide protons of Lys24 and Ala32 that follow the modified Cys^II^ and Cys^III^ residues, respectively, and to the Hα protons of Asp22 and Ser30 that precede them (Fig. 3D). Further analysis of these spectra showed the absence of both Hα and NH protons for the modified Cys residues. The absence of the latter was verified on a natural abundance ^1^H, ^15^N HSQC spectrum (*SI* Fig. S12). These data suggest that their N and Cα atoms are part of heterocycles.

A ^1^H, ^13^C HSQC experiment showed correlations from the Asp22 and Ser30 Hα atoms to their own Cα carbons at 48.1 and 52.4 ppm, respectively, 6 ppm higher than their random coil values (30). The ^1^H, ^13^C HMBC spectrum connected the same Hα protons to carbon atoms at 161.1 and 159.8 ppm, respectively (*SI* Fig. S12). We produced a uniformly ^13^C,^15^N-labeled modified BufA1 molecule in *E. coli* and used its central tryptic peptide to obtain further carbon assignments on the basis of triple-resonance NMR experiments (31, 32). In the HNCACB experiment, we found no resonances that could correspond to upstream neighbors for Lys24 or Ala32, confirming that the Cα carbons of modified Cys^II^ and Cys^III^ do not resonate at the typical 50-ppm value expected for Cys residues (Fig. 3E). In a HNCO experiment starting from the amide groups of Lys24 or Ala32, we found values around 184 ppm for the C=O carbon atoms of both modified Cys residues (Fig. 3F), far from 175-ppm and 200-ppm values expected for such atoms in random-coil and thio-amide peptides, respectively (30, 33). By increasing the resolution and varying the offset of the decoupling pulse during the C=O evolution period, we determined a 90-Hz coupling constant of both carbonyls with carbon atoms whose chemical shift is ∼120 ppm, consistent with their absence in the HNCACB experiment (Fig. 3G). Both parameters point to the C=O carbon atoms of the modified Cys residues connected to carbon atoms in aromatic rings (34).

Comparisons of the chemical shifts assigned on the modified BufA1 peptide with those found for a 5-thiooxazole-containing peptide (28) show the same 6-ppm upfield shift of the Cα carbon of the preceding residue, the presence of similar carbon resonances at 120 and 160 ppm, and the 90-Hz coupling constant of the following carbonyl towards an aromatic ring carbon. Together with the MS data and the characteristic spectrophotometric spectrum (28), we conclude that the Buf1 modification implies cyclization of its central Cys residues to form thiooxazole groups (Fig. 3H).

### Copper binding by bufferin 1

Since heterocycles can be involved in copper binding (35), we interrogated if bufferins can complex with copper, which could be the basis for their protective effect. As attempts to saturate Buf1 with copper *in vitro* were inefficient (*SI* Fig. S13), we investigated complex formation *in vivo* by treating cultures of *bufABCD1*-expressing *E. coli* with CuSO_4_ for three hours prior to bufferin purification. Native MS analysis showed that purified Buf1 thus produced was fully loaded with copper (Fig. 4A). Pulsed electron paramagnetic resonance (EPR) spectra confirmed that Buf1 binds paramagnetic Cu^2+^, and spectral simulations indicated coordinating N atom(s) (36) (Fig. 4B; *SI* Tables S2 and S3).

**Figure 4.**
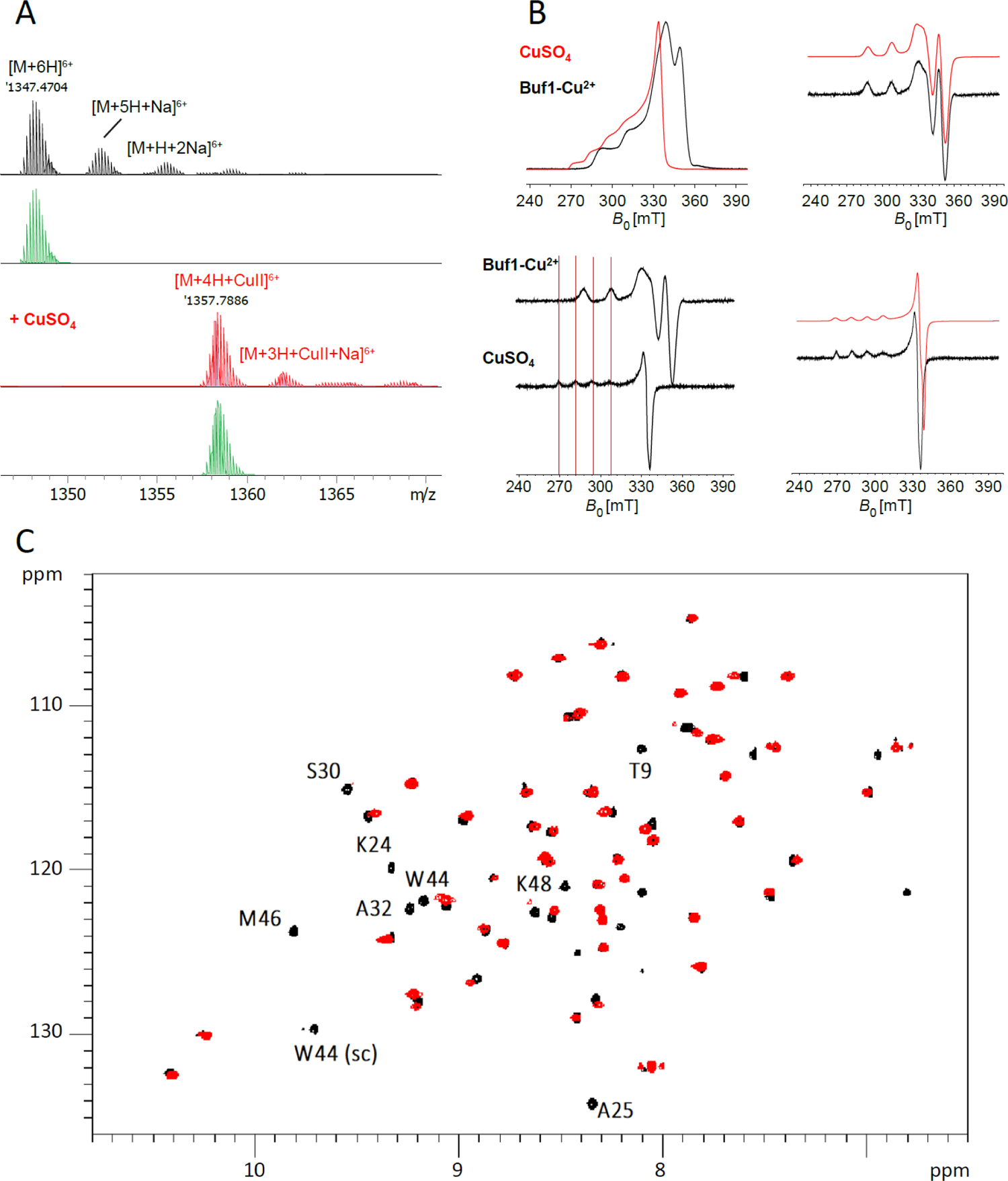
Analyses of the bufferin 1-copper complex formed *in vivo.* **A.** Native MS analysis of the purified complex: isotopic patterns of the major charge state (6+) species. The spectra in black and red show Buf1 from a non-supplemented culture and from cultures supplemented with CuSO_4_, respectively. The calculated isotopic patterns for [M+6H]^6+^ and [M+4H+CuII]^6+^ species are shown in green. **B.** Evidence for Cu^2+^ binding to Buf1 by EPR spectroscopy. Spectra of Echo-Detected Field-Swept of CuSO_4_ (red) and of the Buf1-Cu^2+^ complex (black) are shown with the pseudo-modulation of the spectra underneath. The right panels show the Easyspin fits (red) and the experimental spectra (black) for the Buf1-Cu^2+^ complex (top) and for CuSO_4_ (bottom). **C.** Superposition of the ^1^H, ^15^N HSQC spectra of apo Buf1 (black) and the Buf1-Cu^2+^ complex (red). sc= side chain.

To gain further insight into the binding site, we produced a ^15^N,^13^C-labeled Buf1 sample and recorded triple resonance spectra to assign its resonances. Simultaneously, we used the ^1^H, ^15^N HSQC plane as a basis for HSQC-NOESY and TOCSY experiments to confirm the assignments and gain some knowledge on structural aspects of Buf1. The sequential assignment was interrupted by the modified Cys residues, as expected. Interestingly, amide resonances of the tryptic peptide and of the same segment in the full-length bufferin do not correspond at all, suggesting that the full-length sequence imposes structural constraints on the thiooxazole-containing loop. Part of these likely come from the β sheet spanning two short strands in the structural model, Thr9-Tyr13 and Trp44-Lys48 (*SI* Fig. S1), that we confirmed experimentally through the strong Hα(i-1) – HN(i) NOE contacts between residues in these segments and the cross-strand NOE contacts between the amide protons of Lys45 and Cys12, and of Val47 and Glu10, respectively (*SI* Fig. S14). PTMs of the Cys residues hence do not interfere with the 3D structure predicted for the unmodified sequence.

A comparison of the spectra of the bufferin produced with or without copper showed many peaks unchanged, some experiencing a shift and a signal reduction, and yet others disappearing altogether (Fig. 4C). The latter most probably concern residues close to the paramagnetic Cu^2+^ ion. In particular, disappearance of the peaks corresponding to direct neighbors of the heterocycles, such as Lys24, Ser30 and Ala32, confirms that the thiooxazoles are involved in Cu^2+^ coordination. Interestingly, signals of Thr9, Trp44, Met46 and Lys48 predicted on the β-sheet face oriented towards the thiooxazole-containing loop also disappeared. Our data thus support that the thiooxazole cycles and residues of the β sheet are involved in copper binding.

### BufB1 and BufC1 are required for post-translational modification of bufferin

To dissect the role of individual *buf* genes, we constructed Cv_buf1_ derivatives that lack one or several genes and tested their growth phenotypes in the presence of CuSO_4_. All deletion strains except for that lacking *bufD1* displayed slow-growth phenotypes, showing that BufA1, BufB1 and BufC1, but not BufD1, are required for the function (Fig. 5A).

**Figure 5.**
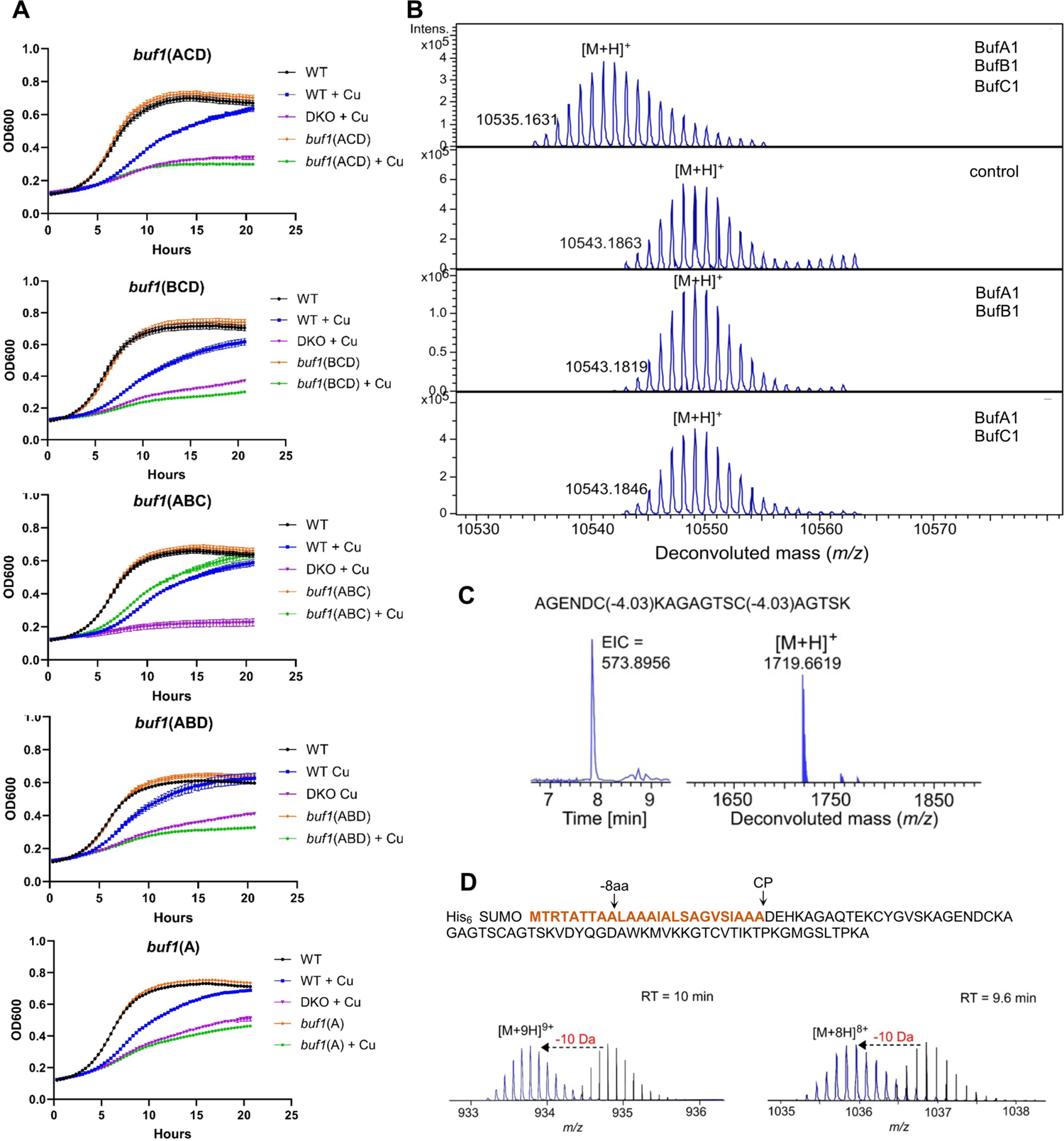
Biosynthesis of bufferin 1. **A.** Effect of Cu on the growth of *C. vibrioides* strains expressing the indicated genes. **B.** *In vitro* reconstitution of Buf1 biosynthesis. The deconvoluted mass spectra for the products of the *in vitro* reactions are shown, with the added proteins indicated in the corresponding panels. Heat-denatured BufB1 and BufC1 were used for the control reaction. **C.** LC-MS analysis of the 19-residue peptide of *in vitro* modified SUMO-Buf1. The extracted ion chromatogram (EIC) of [M+3H]^3+^ (at *m/z* 573.8945) and the deconvoluted mass are shown. **D**. Role of the signal peptide for the PTMs. A schematic representation of BufA1 variants with signal peptide truncations is shown on top. LC-MS analyses of modified BufA1(−8 aa) (left) and modified BufA1(CP) (right): the MS spectra of the most abundant charge states [M+9H]^9+^ and [M+8H]^8+^, corresponding to deconvoluted monoisotopic masses [M+H]^+^ of 8391.1281 and 8277.9398 (blue), respectively, are compared to the theoretical spectra of the unmodified peptides (black).

Since MNIO enzymes require Fe^II^ and Fe^III^ ions for activity (9, 12), purified BufB1 was analyzed by inductively coupled plasma - optical emission spectrometry, showing the presence of two Fe ions. To reveal the role of BufB1 and BufC1 in the bufferin maturation, we reconstituted the modification reaction *in vitro* using the purified proteins (*SI* Fig. S15). LC-MS analyses showed that BufA1 was converted to a species with a −8 Da mass shift only when BufB1 and BufC1 were present (Fig. 5B). Trypsin digestion of the *in vitro* modified BufA1 yielded the 19-residue peptide harboring the thiooxazole groups as confirmed by MS/MS analyses (Fig. 5C). Together with the gene inactivation data, these experiments established that BufB1 and BufC1 are required and sufficient to install the PTMs on BufA1.

### The signal-peptide sequence is dispensable for heterocycle formation

A notable feature of bufferin biosynthesis is that the precursor peptide harbors a Sec signal peptide, which is unprecedented in bacterial RiPPs precursors. We investigated if this sequence also acts as a leader peptide required for Cys transformation. Using the SUMO-*bufA1/bufB1/bufC1 E. coli* co-expression system, the *bufA1* gene was mutated to generate variant precursors with the signal peptide truncated eight residues from the N-terminus (BufA1(−8 aa)) or completely removed (BufA1(CP)). Subsequent LC-MS analysis of purified BufA1 after SUMO-tag cleavage indicated that both truncated precursors were still modified with a −10 Da shift, consistent with the presence of two thiooxazoles and one S-S bond (Fig. 5D). Thus, the signal peptide is dispensable for the installation of the PTMs, suggesting that main recognition determinants of the precursor peptide by the BufB1C1 machinery are located within the core sequence.

## Discussion

Here we report a widespread bacterial strategy for metal homeostasis based on the synthesis of periplasmic RiPP metallophores. Incidentally, we also solved the conundrum of the gold-induced genes found in environmental and pathogenic bacteria (14–16). Bufferins form a new RiPP family with original features and a new function. We showed that they harbor a rare modification, 5-thiooxazole, installed by an MNIO enzyme. With thousands of homologs, bufferins are highly prevalent in eubacteria and represent the largest family of MNIO-modified RiPPs.

The biosynthesis of bufferins involves unique features. To our knowledge they are the first RiPPs naturally exploiting Sec signals for efficient translocation to the periplasm. Peptides with bulky modifications or macrocycles would be poor substrates of the Sec machinery due to the pore size of SecY (37). Although bufferins harbor an S-S bond formed macrocycle, the latter is most likely installed in the oxidizing periplasmic environment after export. The use of ‘generalist’ systems, Sec and signal-peptidase, for export and proteolytic maturation is a new paradigm in the RiPP field.

The precursors and mature forms of bufferins are large compared to those of other MNIO-modified RiPPs, and our *in-silico* analyses identified yet larger bufferin-like precursors of more than 200 residues with numerous Cys residues. Thus, the bufferin family appears to harbor a continuum from modified peptides to proteins, which expands the concept of *bona fide* RiPPs to what could be ‘RiPProteins’.

The conversion of Cys to 5-thiooxazole represents a new reaction catalyzed by MNIOs, further highlighting their chemical versatility. Intense efforts have been made recently to explore the chemical space of RiPPs undergoing MNIO-catalyzed transformations (7, 8, 11, 13, 38). In line with previous work on Cys-modifying MNIO enzymes, BufB would similarly make use of the mixed-valent iron center and proceed by an intermediate step of proton abstraction from the Cβ of cysteine (9, 10, 12, 13) (SI Fig. S16).

We demonstrated that model bufferins are involved in copper homeostasis by complexing with the metal in the periplasm. This novel function for RiPPs contrasts with that of other MNIO-modified metallophores, methanobactins, a phylogenetically restricted family of RiPPs which essentially scavenge copper ions in the extracellular milieu for its provision to cuproprotein clients (39–42). The mode of binding of bufferins also differs from that of methanobactins. Notably, *in vivo* formation of the bufferin-Cu^2+^ complex most likely occurs while the unfolded bufferin emerges from the Sec machinery in the periplasm and folds around copper, which explains that our *in vitro* loading attempts were not successful. The defensive role of bufferins is reminiscent of those of non-ribosomal peptides, anthrochelin and yersiniabactin (43, 44), although the putative multiple metal binding sites of large bufferins suggest that some of them might serve as intracellular metal reservoirs. Future studies on this large, diverse family will likely also unveil roles for bufferins in the homeostasis of other transition metals.

Copper is a major metal pollutant from anthropic sources (45, 46), and bufferins represent a previously unknown but highly prevalent mechanism of bacterial adaptation likely relevant in the environment, agriculture and healthcare. In *C. vibrioides*, their expression is induced at high copper concentrations, suggesting that they come into play after other defense systems are overwhelmed. Genetic determinants that provide environmental species with protection against transition metals can contribute to the emergence of opportunistic pathogens (47), as exemplified by *Legionella pneumophila* which harbors up to six bufferin BGCs. Bufferins are most likely beneficial to bacteria in host-pathogen interactions notably in phagosomal compartments, which might account for their presence in *Bordetella pertussis, Neisseria gonorrheae, Haemophilus influenzae* and *Pseudomonas aeruginosa* (26, 48, 49). From a more applied perspective, it should be possible in the long term to engineer bufferin variants for bioremediation purposes (3).

## Materials and methods

### Genetic constructs and growth conditions

Construction details, plasmids, recombinant strains, primers and synthetic genes are described in *SI* Text and Tables S4 to S6, respectively.

### Functional experiments

*C. vibrioides* growth curves were recorded using a TECAN Spark plate reader, with OD_600_ measurements every 20 min. The gain-of-function experiment was performed in *E. coli* expressing the *buf*1 operon under the control of an IPTG-inducible promoter. The bacteria were challenged with 0.3 mM CuSO_4_ for 4 hours. For the predation experiments, *Dictyostelium discoidum* was placed in the center of bacterial lawns before incubation at 20°C for three days, and the densities of bacteria in the contact zones were determined with ImageJ. Experimental details are in *SI* Text.

### Bufferin production and purification

For Buf1 or Buf2 production in *C. vibrioides*, strains harboring the pSigF plasmid were grown overnight in PYE medium containing 100 μM IPTG, and the bufferins were purified on Streptactin or Ni-NTA columns using standard procedures. Production of isotopically labeled Buf1 was performed in *E. coli* BL21. For the *in vitro* assays, N-His_6_-SUMO-BufA1, N-His_6_-BufB1 and N-His_6_-BufC1 were produced in BL21(DE3), purified on Ni-NTA columns, with 10 mM Tris(2-carboxyethyl)phosphine hydrochloride added to all buffers for BufA1. Production of the trypsic peptide for NMR studies was performed in BL21(DE3) co-expressing Sumo-BufA1, BufB1 and BufC1. The iron content of His_6_-BufB1 was measured using an inductively coupled plasma - optical emission spectrometer (ICP-OES 5110 VDV, Agilent Technologies). Experimental details are provided in *SI* Text.

### Peptide analyses by MS

All experimental details are given as *SI* text.

### NMR analyses

The central tryptic peptide was purified by preparative HPLC. All NMR spectra were recorded at 293 K on an 800-MHz NEO Bruker spectrometer equipped with a QCP cryogenic probe head. Experimental details are in *SI* Text.

### EPR experiments

Pulsed EPR experiments were performed on the Buf1^twstr^-copper complex using an ELEXSYS E-580 spectrometer (Bruker). Experimental details are in *SI* Text. Numerical simulation of the spectra was conducted using the Matlab toolbox Easyspin 5.2.35.

### Spectrophotometric copper binding assay

This was performed as described (50).

### Enzymatic assays in vitro

Purified His-SUMO-BufA1 was digested with SUMO protease before adding either or both N-His_6_-BufB1 and N-His_6_-BufC1. After incubation the mixtures were analyzed by LC-MSMS. Details are presented as *SI* Text.

### In silico analyses

The search for putative bufferin precursors and the SSN analyses are detailed in *SI* Text.

## Acknowledgments

We thank E. Lesne and G. Roy for initiating this work long ago, J.-M. Saliou for preliminary MS analyses and J.-Y. Matroule and P. Cherry for providing *C. vibrioides* strains, plasmids and protocols. This project was funded by the ANR grant CuRiPP (ANR-22-CE44-0001-02) to FJD. L. Leprevost and S. Jünger were supported by PhD fellowships of Lille University and the French Ministry of Education (ED 227 MNHN-SU), respectively. S. Dubiley was supported as a Visiting Scientist by a fellowship of the Collège de France. We thank the NMR facility of MetaToul (www.metatoul.fr). Metatoul is part of the French National Infrastructure for Metabolomics and Fluxomics MetaboHUB-AR-11-INBS-0010 (www.metabohub.fr), and is supported by the Région Midi-Pyrénées, the ERDF, the SICOVAL and the French Minister of Education & Research, who are all gratefully acknowledged. The LC-MS data were acquired at the MNHN bioorganic mass spectrometry platform and the PAPPSO platform (http://pappso.inra.fr/en). For the EPR experiments the financial support from the IR INFRANALYTICS FR2054 is acknowledged. ICP-OES measurements were performed by V. Alaimo on the Chevreul Institute Platform (U-Lille / CNRS). The Region Hauts de France and the French government are acknowledged for funding this apparatus.

