## Supplementary Text, Figures and Tables for "A widespread family of ribosomal peptide metallophores involved in bacterial adaptation to metal stress"

| **This pdf file includes** | Supporting text |
|---|---|
| | Figures S1 to S16 |
| | Tables S1 to S6 |
| | SI References |

### Supporting text

### Materials and Methods

***C. vibrioides constructs and growth conditions.*** Strain NA1000 was used. Allelic exchange was performed using pNPTS138. pSRK-Km was used for expression of SigF. Plasmids were mobilized from *E. coli* S17 into *C. vibrioides* by conjugation. Expected deletions were verified by PCR. Chromosomal transcriptional fusions were generated using pFUS2 (1). Plasmids, recombinant strains, primers and synthetic genes are shown in Tables S4 to S6. *C. vibrioides* was grown at 30°C in PYE medium with nalidixic acid (30 and 15 µg/ml in solid and liquid medium, respectively), and kanamycin (20 and 5 µg/ml) or gentamycin (5 and 0.5 µg/ml) when carrying specific plasmids. To record growth curves, cultures in the stationary phase were diluted to $OD_{600}$ = 0.05 and inoculated in 96-well plates with continuous shaking at 30°C in a TECAN Spark plate reader, with $OD_{600}$ measurements every 20 min for 28 h. $CuSO_4$ was added to 225 µM where indicated. Reporter strains were grown to early stationary phase in the presence of metal ions at the indicated concentrations, and β–Gal activities were measured as described (1). Biological quadruplicates were obtained in all assays.

***Genetic constructs in E. coli.*** For expression in *E. coli*, the *buf1* operon was PCR-amplified and inserted after the T5-lac promoter of pCA24 (2) using Gibson assembly (3). The PCR-amplified sequence coding for BufA1 with a C-terminal Strep-tag II was used to replace the native *bufA1* gene, yielding pCA24-A$^{str}$BCD. For large-scale production of recombinant BufA1, the PCR-amplified gene was inserted into SspI-linearized pETHisSUMO by ligase-independent cloning, yielding pETHisSUMO-A1. pACYC-Duet-based expression plasmids were created using *in vivo* assembly (IVA) (4) yielding pACYC::HisC1 *and* pACYC::HisB1. For co-expression, *bufB1* was cloned into the MCS2 of pACYC::HisC1 by IVA cloning, yielding pACYC::HisC1B1Stag.

***Gain-of function in E. coli.*** BL21(pCA24-*bufA1*$^{str}$*1BCD*) and BL21(pCA24-*psmCA*) (5) were grown overnight (o/n) at 37°C, diluted 100-fold in M9 medium containing 1xBME vitamin solution (Sigma), 0.5% glycerol, 34 µg/mL chloramphenicol and 1 mM IPTG and grown at 37°C with vigorous shaking for 4 h. At $OD_{600}$ = 0.1, one half of each culture was exposed to 0.3 mM $CuSO_4$. Incubation was pursued for 3 h before CFUs were counted.

***Amoeba experiments.*** Exponentially growing *C. vibrioides* cultures were concentrated to $OD_{600}$ = 1 in PYE medium and plated on PYE-agar with 1% glucose. 50-µL drops of *Dictyostelium discoidum* grown to 80% confluency in HL/5 medium at 20°C and diluted to $10^6$/mL were placed on the plate center, before incubation at 20˚C for three days. ImageJ was used to measure bacterial densities in the zones of lysis, divided by the total areas of these zones.

***Native bufferin production and purification*.** Cultures of $Cv_{buf1}^{TwStrep}$(pSigF), $Cv_{buf1}^{6His}$(pSigF) and $Cv_{buf2}^{6His}$(pSigF) were started at $OD_{600}$ = 0.1 in PYE with 100 µM IPTG. Bacteria were harvested after o/n growth, resuspended in 100 mM Tris-HCl (pH = 8), 150 mM NaCl and broken using a French press. $Buf1^{TwStrep}$ was purified from the clarified lysate using a 1-mL StrepTrap XT column (Cytiva) following the manufacturer's instructions, concentrated using Amicon ultrafiltration centrifugal units and dialyzed against 25 mM Hepes (pH 8) 150 mM NaCl. The $Buf1^{6His}$ and $Buf2^{6His}$ proteins were purified on 1-mL $Ni^{2+}$ columns using standard procedures. For purification of isotopically labeled Buf1, *E. coli* BL21(pCA-$A^{str}$BCD) was grown in M9 medium with 34 µg/mL chloramphenicol, 1xBME vitamin solution, 0.5% glycerol-$^{13}C_3$ and 0.1 g/L $^{15}NH_4Cl$ at 37 °C until OD600 ~ 0.6. Protein expression was induced with 1 mM IPTG, and the culture continued at 22°C o/n. $CuSO_4$ was then added to 0.3 mM. After 4 more hours bacteria were harvested by centrifugation, resuspended in 50 mM Tris-HCl buffer (pH 8.0), 300 mM NaCl and lysed by sonication. The clarified lysate was applied to a 1-mL Strep-Tactin® Superflow column (IBA Lifesciences, Germany), and after washing with 20 column volumes, $Buf1^{str}$ was eluted with 2.5 mM desthiobiotin, 50 mM Na-phosphate buffer (pH 7.0), 50 mM NaCl. The molar extinction coefficient of $Buf1^{str}$ is 38,158 $M^{-1}cm^{-1}$.

***Protein production and purification for in vitro assays.*** N-$His_6$-BufB1 and N-$His_6$-BufC1 were each produced in *E. coli* BL21(DE3), in 1 L autoinduction ZYM-5052 medium (6) containing 250 µM ferrous ammonium sulfate for BufB1. After 4 h at 37°C, the temperature was lowered to 18°C and cultures were pursued o/n. N-$His_6$-SUMO-BufA1 was produced in 1 L TB medium, with bacteria grown until $OD_{600}$ = 0.6-0.7 before cooling down on ice. After adding IPTG to 0.1 mM, the cultures were pursued at 18°C o/n. Bacteria were resuspended in 50 mM Tris-HCl (pH 7.6), 300 mM NaCl, 10% glycerol, 10 mM imidazole and lysed using

French Press or by sonication in the case of BufA1. Recombinant proteins were purified by gravity using a self-packed column with 5 mL of Ni-NTA resin slurry (Qiagen). After washes with 30 mM and 60 mM imidazole in 50 mM Tris-HCl (pH 7.6), 300 mM NaCl, the proteins were eluted with 50 mM Tris-HCl (pH 7.6), 300 mM NaCl, 250 mM imidazole, concentrated by ultrafiltration and buffer exchanged by PD10 columns (Cytiva) to 50 mM Tris-HCl (pH 7.6), 150 mM NaCl, 20% glycerol. Purified proteins were flash frozen and stored at -80°C. Purification followed the same procedure for all three proteins except that 10 mM of Tris(2-carboxyethyl)phosphine hydrochloride (TCEP) was added to all buffers for BufA1. The iron content of $His_6$-BufB1 was measured by using an inductively coupled plasma - optical emission spectrometer (ICP-OES 5110 VDV, Agilent Technologies) at $\lambda$ = 238.204 nm calibrated using standard solutions. All solutions were acidified with $HNO_3$ (Fisher Scientific, 67–69%, Optima and trace metal grade). Certified water Enviromat (SCP, N°140-025-038) was analyzed for quality control (certified value of 0.581 +/- 0.011 mg/L, experimental result of 0.59 +/- 0.02 mg/L). The limit of detection was estimated to 0.005 mg/L and the average blank value was below this limit.

***Production of SUMO-BufA1 for NMR studies.*** *E. coli* BL21(DE3, pACYC::HisC1B1Stag, pETHisSUMO-A1) was grown in 2 L TB medium. For $^{13}C$, $^{15}N$-labeling, protein expression was carried out in 1 L M9 medium containing 1 g/L of $^{13}C$-glucose and 5 g/L $^{15}NH_4Cl$. Cultures were grown until $OD_{600}$ = 0.6-0.7 before cooling down on ice, and after adding ITPG to 0.1 mM they were pursued at 18°C for 16 h (TB medium) or 48 h (M9 medium), respectively. Protein purification was performed as described above.

***Peptide analyses by MS.*** Cell pellets and culture supernatants from 25-mL o/n *C. vibrioides* cultures were resuspended in 1 mL 50 mM Tris-HCl (pH 7.5) and submitted to fast-prep lysis in Lysing Matrix B tubes (force 5, 30 sec, 2 times). The clarified lysates were submitted to solid-phase extraction (SPE) on Sep-Pak tC2 Plus Light Cartridges (Waters) conditioned with 5 mL methanol and 5 mL 0.01 % formic acid (FA). The cartridges were washed with 5 mL 0.01 % FA and eluted with 2 mL methanol. The eluted fractions were split in two, evaporated under vacuum and dissolved in 50 µL 0.01% FA/methanol 1:1 (v/v). One was left intact (I) and the other treated with 10 mM dithiothreitol (DTT) for 1 h at 50°C and alkylated with 15 mM iodoacetamide for 1 h at 20°C in the dark. Half of that solution was collected (R1) and the other half was treated with 0.01 µg/µL trypsin (Gold, Promega) o/n at 37°C (R2). Alternatively, reduction-alkylation was performed using DTT together with 2-bromoethylamine (7). R1 and R2 were desalted using Pierce C18 Spin columns. Purified Buf1$^{twstr}$ was analyzed by LC-MS directly or after reduction-alkylation and digestion with trypsin as above. I, R1 and R2 were analyzed by ultra-high-performance liquid chromatography-MS on an Ultimate 3000-RSLC system (Thermo Scientific) connected to an electrospray ionization- quadrupole–time of flight instrument (Maxis II ETD, Bruker Daltonics). The separation was achieved using an Aeris widepore C4 column (3.6 µm, 200 Å, 2.1 × 100 mm, Phenomenex) for I and R1 samples and a RSLC Polar Advantage II Acclaim column (2.2 µm, 120 Å, 2.1 × 100 mm, Thermo Scientific) for R2, with a linear gradient (2-80 % in 15 min) of mobile phases FA 0.1% and LC-MS grade acetonitrile (ACN) + FA 0.08%, at 300 µL/min. For I and R1, LC-MS detection was carried out in positive mode in the *m/z* range 250-2500. For R2 samples, data-dependent LCMS/MS acquisitions were conducted. They were also analyzed by nanoLC-MS on an UltiMate 3000 RSLCnano System coupled to an Orbitrap Fusion Lumos Tribrid. The separation was achieved using a C18 Biosphere column (2 µm, 75 µm × 500 mm Neo Thermoscientific) with a linear gradient (2.5-45% B for 55 min, 45-98% for 10 min and 2.5% for 10 min) of phases A (FA 0.1%/ACN 98/2) and B (FA 0.1%/ACN 20/80) at 0.250 µL/min. The mass spectrometer was operated in positive mode in the *m/z* range 150-1200. Data-dependent LCMS/MS acquisition mode was used. Full MS scans were captured with a resolution of 120,000. Peptide identification was performed using X!TandemPipeline (8) or PEAKS Studio X Pro (Bioinfomatics Solutions Inc.), against the *Caulobacter* UniProtKB database (version 2023) with the following variable modifications: one possible missed cleavage, Cys carboxyamidomethylation, and Met oxidation. Precursor mass and fragment mass tolerance were 10 ppm and 0.02 Da, respectively. Data filtering was achieved according to a peptide E value < 0.05. The false discovery rate at the peptide level was assessed from searches against a decoy database. MS2 spectra were analyzed using BYONIC™ (Protein Metrics) together with i2MassChroQ 0.6.7 (http://pappso.inrae.fr/bioinfo/i2masschroq) and SpecGlobTool to highlight and localize unknown PTMs (9). Native MS was carried out using flow injection with 50 µL/min 10 mM ammonium acetate (pH 7). One µL of purified peptide at 20 µM in the same buffer was injected.

***NMR analyses.*** Purified His-SUMO-BufA1* was incubated with trypsin (1:100 w/w) at 37°C o/n in 50 mM Tris-HCl (pH 7.6), 10 mM TCEP. Purification of the 19-residue central trypsin fragment was conducted by HPLC (UltiMate™ 3000 System) on a preparative C18 column (Luna C18 (2), 5 µm, 100 Å, 250 x 10 mm, Phenomenex) at room temperature. Separation was performed using a linear gradient (6 min at 5% B, 6-14 min from 5-30 % B, 14-18 min to 80% B, and 18-22 min at 80%B) of phases A (0.1% FA) and B (ACN) at 3 mL/min and monitored at 280 and 305 nm. Fractions containing the relevant peptide were pooled and dried by SpeedVac. A sample of 3.4 mg of the peptide dissolved in 550 µL of 50 mM Na phosphate buffer (pH 6.5), 150 mM NaCl, with 25 µL $D_2O$ and Trimethyl Silyl Propionate (TSP) as reference was introduced in a standard 5-mm NMR tube. All spectra were recorded at 293 K on an 800-MHz NEO Bruker spectrometer equipped with a QCP cryogenic probe head. TOCSY and ROESY spectra were recorded for assignment of the peptide with 16k x 512 points for a spectral width of 12 ppm in both dimensions, with 16 and 64 scans per increment, respectively (total measurement times of 4 h and 16 h, respectively). Scalar coupling transfer was accomplished during a 60-ms DIPSI2 mixing time, whereas a 100-ms spin lock was used for the dipolar magnetization transfer. The relaxation delay was set to 1 s for both spectra. The natural abundance $^{1}$H, $^{15}$N HSQC spectrum was recorded with the Bruker hsqcf3gpph19 pulse program, with 2k x 128 points for a spectral width of 12 ppm x 36 ppm, centered on 4.7 ppm and 119 ppm for the H and the N dimensions, respectively, and 256 scans per increment. Natural abundance $^{1}$H, $^{13}$C HSQC and HMBC spectra were recorded with standard Bruker pulse programs (hsqcetgpsisp.2 and hmbcedetgpl3nd), with 2k x 256 points for a spectral window of 12 x 40 ppm and the $^{13}$C carrier set at 80 ppm, and with 2k x 256 points for a spectral window of 12 x 140 ppm and the $^{13}$C window centered at 100 ppm, respectively, and 64 scans per increment for both. A HNCACB spectrum was recorded on the $^{15}$N, $^{13}$C labeled peptide as a cube of 12 x 30 x 70 ppm sampled with 2k x 76 x 256 points centered at 4.7 x 119 x 40 ppm in the $^{1}$H, $^{15}$N and $^{13}$C dimensions, respectively. For the HNCO spectrum, the $^{13}$C carrier was shifted to 175 ppm, and the $^{13}$CO window of 30 ppm was sampled with 64 points. In order to estimate the frequency of the carbon coupled to the carbonyl of the modified Cys residues, we recorded the H(N)CO planes of the latter spectrum while introducing during the $^{13}$CO evolution period a selective pulse with varying offset instead of the classical C$\alpha$ selective 180° pulse.

***EPR experiments.*** Pulsed (EDFS) EPR experiments on the *in vivo*-formed Buf1$^{twstr}$-$Cu^{2+}$ complex (200 μM) were carried out using an ELEXSYS E-580 spectrometer (Bruker) equipped with a Super-Q FTu bridge for X- (9 GHz) and Q- (35 GHz) band experiments. Samples were drawn into 3 × 4 mm quartz tubes, flash-frozen into liquid nitrogen and rapidly transferred into precooled dielectric resonators, ER 4118X-MD5W X-band and ER 5106QT-2w Q-band. Both X- and Q-band spectra were acquired at 20 K using a Bruker Cryogen-free (Cold-Edge) cooling system. Microwave pulse lengths $t_{\pi/2} = 10\text{-}12$ ns (Q-band) and $t_{\pi/2} = 16$ ns (X-band) were used with inter-pulse delays of $\tau = 172$ ns (Q-band) and $\tau = 132$ ns (X-band). A two-step phase-cycle was applied to remove all unwanted echoes on EDFS. Numerical simulation of the EPR spectra was conducted using the Matlab toolbox Easyspin 5.2.35.

***Spectrophotometric copper binding assay***. Purified $Bufl^{twstr}$ (5 to 40 µM) was used in competition with a complex between 4-(2-pyridylazo)resorcinol (PAR) and $Cu^{2+}$ ($\varepsilon_{505nm}$ =41,500 $M^{-1}$ $cm^{-1}$) (4 µM Cu and 10 µM PAR) (10). UV/visible spectra were recorded after increasing incubation periods at room temperature.

***Enzymatic assays in vitro***. Reactions were carried out in 100-µl solutions of 50 µM purified His-SUMO-BufA1 in 50 mM Tris-HCl (pH 7.6), 10 mM TCEP. This mixture was first incubated with in-house purified SUMO protease at a 1:20 ratio of protease/peptide o/n at 4°C. Subsequently, either or both purified N-$His_6$-BufB1 and N-$His_6$-BufC1 (25 µM each) were added, and the mixtures were incubated at 30°C for 16 h. For control reactions BufB1 and BufC1 were inactivated by heating. Half of the reaction mixture was desalted using Zeba spin columns (Thermo Fisher Scientific) and directly analyzed by LC-MSMS on a Aeris widepore C4 column. The other half was treated with trypsin added to 1:100 (w/w) and incubated for 3 h at 37°C before LC-MSMS analysis on a RSLC Polar Advantage II Acclaim column. LC-MSMS measurements were performed on an Ultimate 3000-RSLC system connected to a high-resolution ESI-Q-TOF mass spectrometer (Maxis II ETD), as above.

***In silico analyses.*** The non-redundant NCBI database (release of Jan 2023) was searched for MNIO-coding genes. A script was designed to collect genes next to and in the same orientation as MNIO genes, coding for proteins less than 200 residues long with Cys residues and a signal peptide. Predictions of signal peptides were performed using Scotopus, Philius, Octopius, Scampi, Polyphobius and Topcons. Analyses by the Enzyme Function Initiative - Enzyme Similarity Tool were carried out with the predicted mature proteins to build sequence similarity networks. To generate the BUF_6/12Cys hmm profile, the procedure described in (11) was followed. Briefly, a few dozen proteins were picked from the cluster of interest of the Representative Node Network, and their sequences were aligned with MAFFT (L-INS-i). After manual editing of the non-conserved N- and C-terminal regions, a hmm profile was generated using this seed alignment.

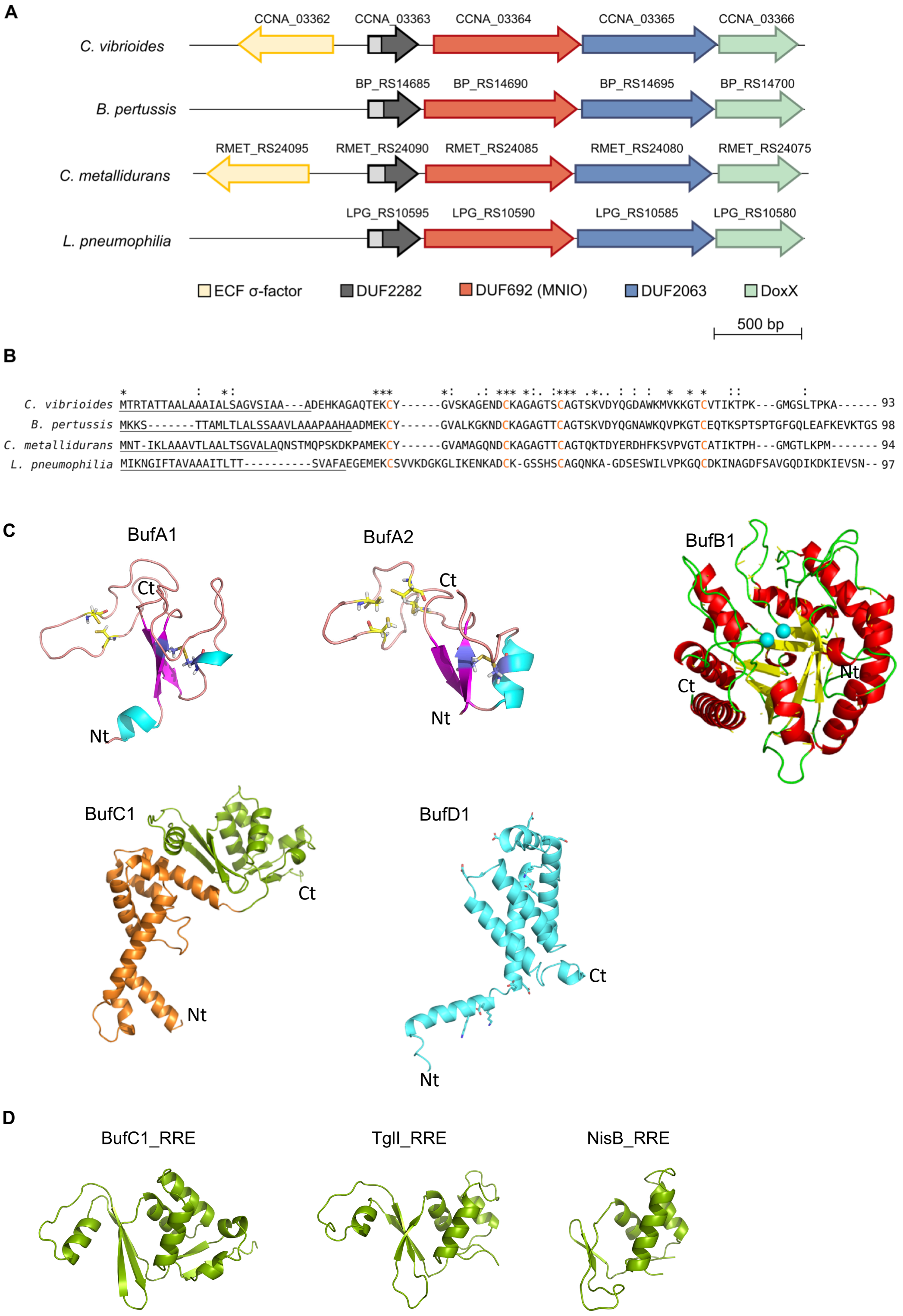


**Figure S1. Main features of the *buf* BGCs and proteins**. **A**. Gene organization of *gig*-like operons found in the genomes of *C. vibrioides* (NC_011916.1), *B. pertussis* (NC_002929.2), *C. metallidurans* (NC_007974.2) and *L. pneumophilia* (NC_002942.5). Signal peptide-coding sequences are shown in pale grey. **B**. Sequence alignment of DUF2282 proteins from *gig*-like operons. Predicted signal peptides are underlined, and conserved Cys residues are highlighted in orange. Sequence alignment was done with Clustal Omega, and signal peptides were predicted using the SignalP 5.0 online tool. **C**. AlphaFold2 models for Buf1 and Buf2 (core peptides, after signal peptide removal), BufB1, BufC1 and BufD1(DoxX) proteins of *C. vibrioides*. The BufC1 protein contains an N-terminal all-α-helical DUF2063 domain (orange) and a C-terminal RRE-like domain (green). In DoxX the 4-helix bundle is predicted to be inserted in the cytoplasmic membrane. Nt and Ct show the N and C termini of the proteins. **D**. Comparisons of the structural model of the C-terminal domain of BufC1 with the crystal structures of the RRE domains of NisB (pdb ID 4WD9) and TglI (pdb ID 8HC1) showing their structural relatedness.

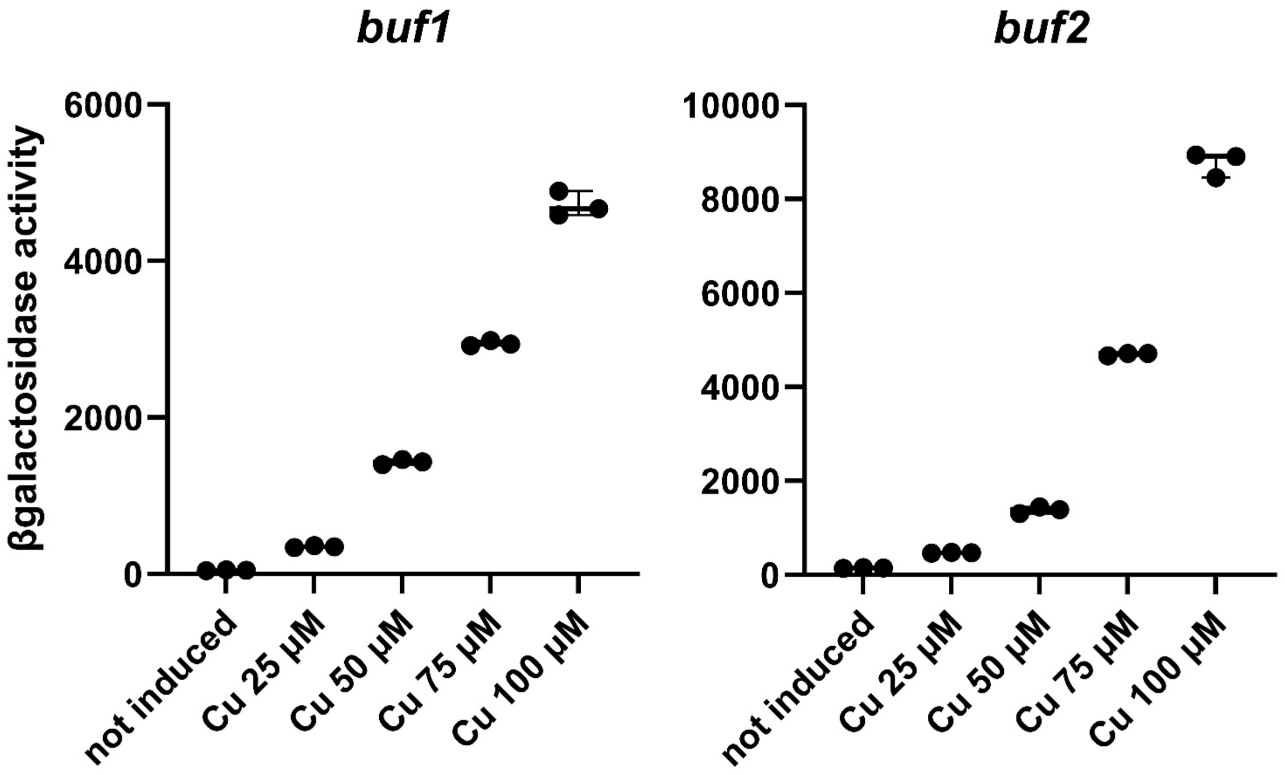


**Figure S2. Reporter assays with *bufA1-lacZ* and *bufA2- lacZ* transcriptional fusions.** Bacteria were grown for 16 h with the indicated concentrations of $CuSO_4$. These experiments were performed in biological triplicates, with three technical replicates.

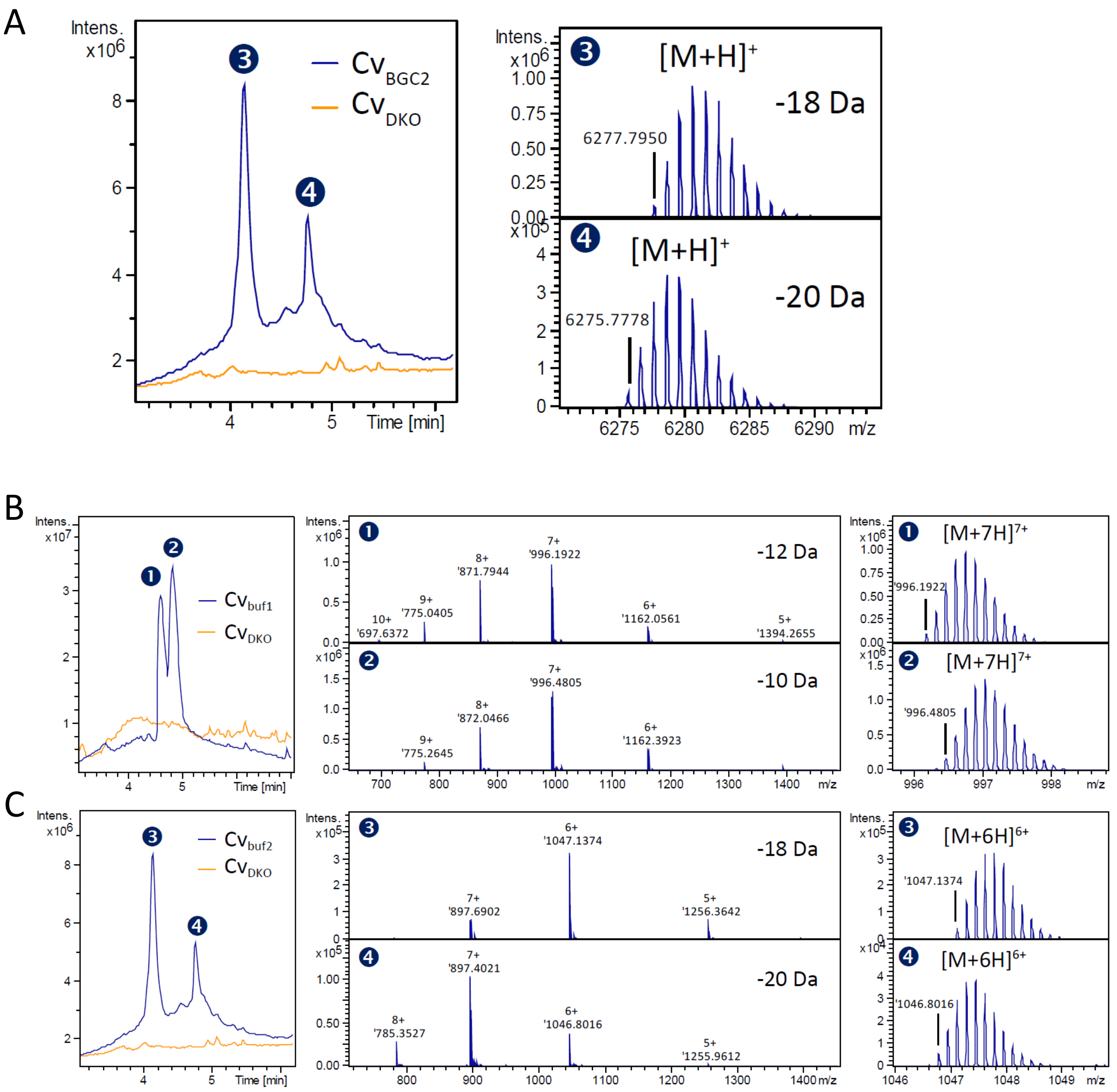


**Figure S3. Top-down analyses of bufferins.** A. LC-MS analyses of the cell extracts of $Cv_{\mathit{buf2}}$(pSigF) compared with $Cv_{DKO}$(pSigF). Left panel: total ion chromatogram, right panels: deconvoluted mass spectra of the two main peaks detected only for $Cv_{\mathit{buf2}}$. Two compounds were detected with masses corresponding to the expected core peptide (after removal of the signal peptide; calculated monoisotopic Mw 6294.96 Da), with mass shifts of -18 Da ($[M+H]^+$ at *m/z* 6277.79) and -20 Da ($[M+H]^+$ at *m/z* 6275.78). B and C. Raw spectra of the LC-MS analyses of the cell extracts of $Cv_{\mathit{buf1}}$(pSigF) (B) or $Cv_{buf2}$(pSigF) (C). Left panels: total ion chromatograms compared to those of the non-producing strain $Cv_{DKO}$(pSigF), middle panels: raw spectra corresponding to the deconvoluted spectra shown in Fig. 2A and in Fig. S3A, respectively, and right panels: zooms on the isotopic patterns of a single multiprotonated species, $[M+nH]^{n+}$.

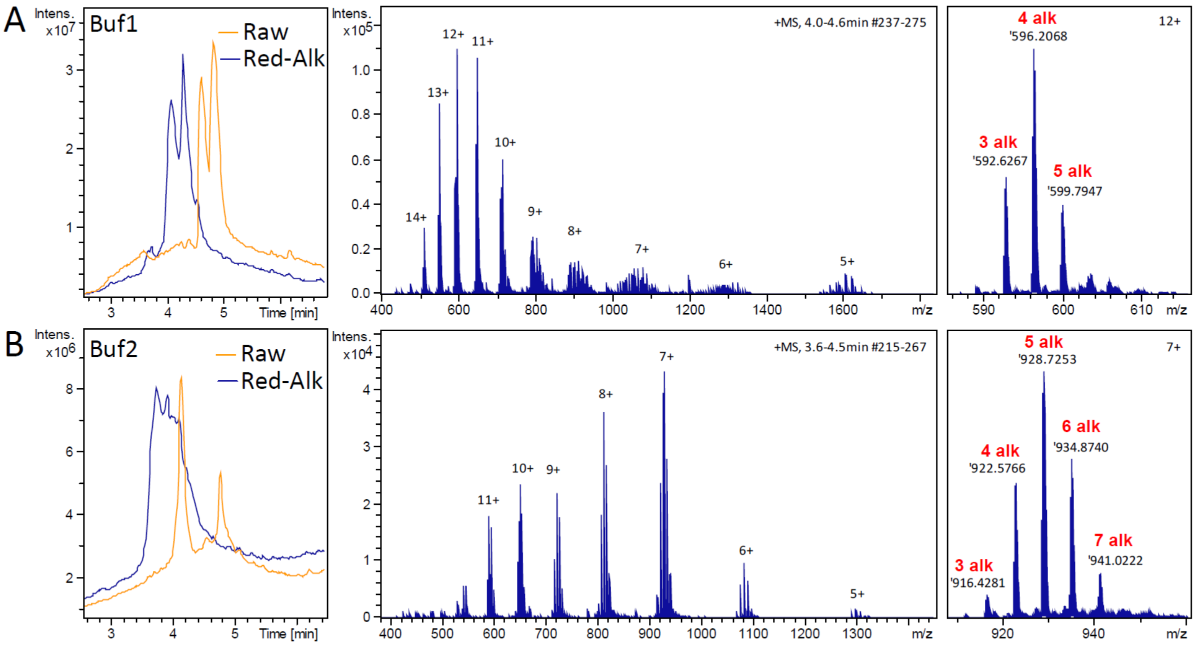


**Figure S4. Reduction-alkylation of Buf1 and Buf2.** Top-down LC-MS analyses of the cell extracts of $Cv_{buf1}$(pSigF) (A) or $Cv_{buf2}$(pSigF) (B) submitted to reduction and alkylation with 2-bromoethylamine. Total ion chromatograms compared to that of the untreated extracts (left panels), MS spectra of the main alkylated species detected (middle panels) and zooms on the major charge states (right panels). These data indicate that residues other than Cys are modified or the PTMs of the Cys residues result in alkylation-sensitive groups. Minor species corresponding to additional unspecific alkylations (5 and 7 alkylations for Buf1 and Buf2, respectively) were also detected.

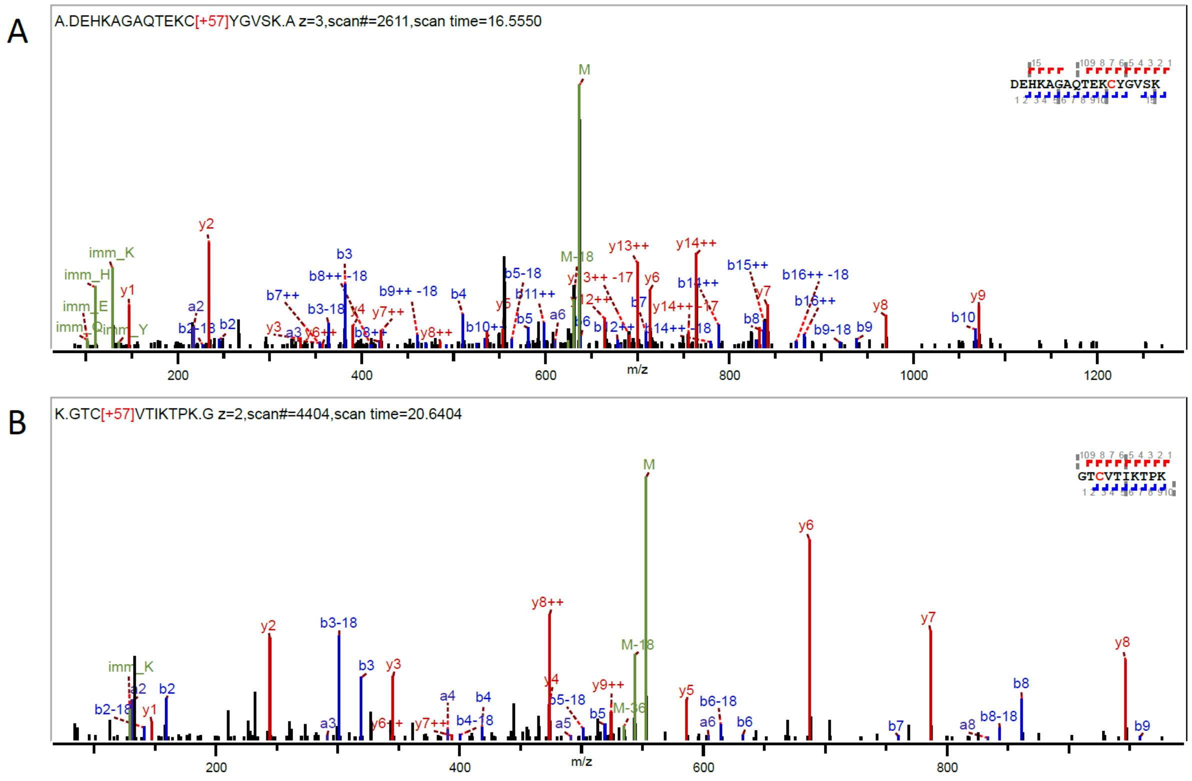


**Figure S5. Bottom-up analysis of bufferin 1.** MS/MS spectra of the tryptic peptides containing the $Cys^{I}$ and $Cys^{IV}$ residues, $^{1}$DEHKAGAQTEKCYGVS$^{17}$K ($[M+3H]^{3+}$ at *m/z* 636.63) (A) and $^{50}$GTCVTIKTP$^{59}$K ($[M+2H]^{2+}$ at *m/z* 552.81) (B), respectively. The two Cys residues carry each a carbamidomethyl group resulting from alkylation with iodoacetamide (+ 57.02 Da).

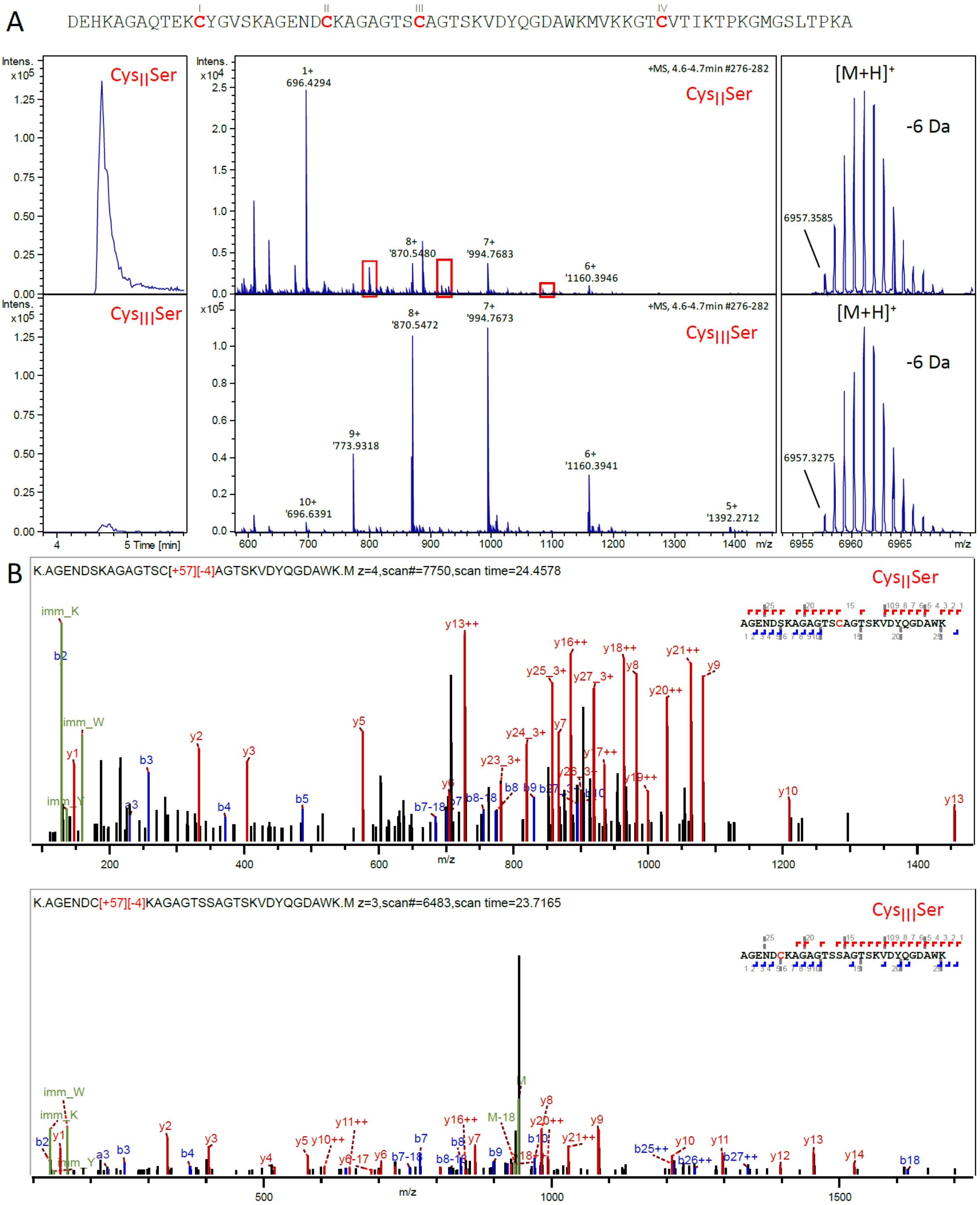


**Figure S6. Analysis of the Cys$^{II}$Ser and Cys$^{III}$Ser variants of Buf1.** Top-down LC-MS analyses of the cell extracts of Cv$_{buf1}$$^{CysIISer}$(pSigF) and Cv$_{buf1}$$^{CysIIISer}$(pSigF). A. Extracted ion chromatograms of the $[M+7H]^{7+}$ species of Buf1$^{CysIISer}$ or Buf1$^{CysIIISer}$ (*m/z* 995) (left panels), corresponding mass spectra (middle panels) and deconvoluted spectra (right panels). The measured *m/z* values (deconvoluted $[M+H]^+$ species at *m/z* 6957.36 and 6957.33 for Buf1$^{CysIISer}$ and Buf1$^{CysIIISer}$, respectively) show -6 Da mass shifts from the masses expected for the proteins with no PTM (calculated monoisotopic Mw 6962.376951). This most likely corresponds to an S-S bond between Cys$^{I}$ and Cys$^{IV}$, which would be consistent with the AlphaFold2 model, and a -4 Da mass shift for the remaining modified Cys$^{II}$ or Cys$^{III}$ residue. B. MS/MS spectra of the tryptic peptides $^{18}$AGEND**S**KAGAGTS**C**AGTSKVDYQGDAW$^{45}$K for Buf1$^{CysIISer}$ ($[M+4H]^{4+}$ at *m/z* 707.56) and $^{18}$AGEND**C**KAGAGTS**S**AGTSKVDYQGDAW$^{45}$K for Buf1$^{CysIIISer}$ ($[M+3H]^{3+}$ at *m/z* 943.08). For both variants, the unsubstituted Cys residues carry a carbamidomethyl group resulting from alkylation with iodoacetamide (+ 57.02 Da), together with a -4.03 Da mass shift.

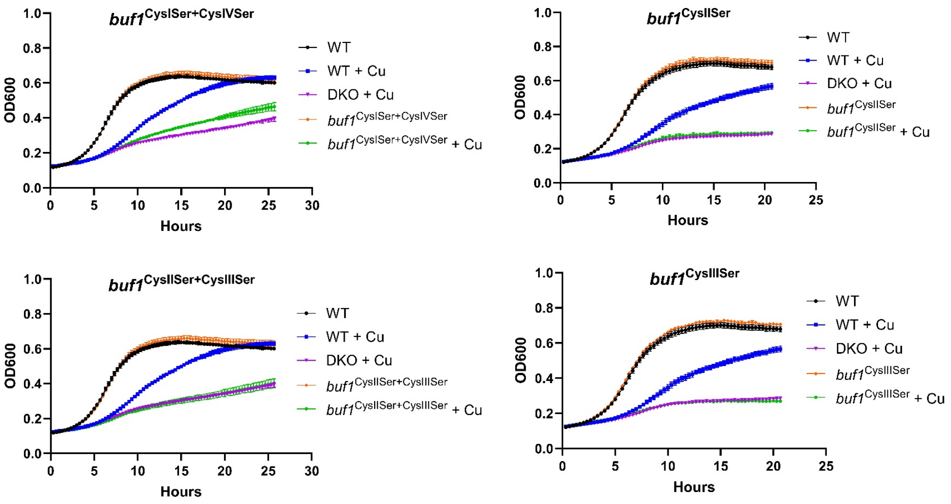


**Figure S7. Effect of Cu on the growth of *C. vibrioides* harboring Cys to Ser substitutions in BufA1**. Strains expressing the *buf1* BGC with the indicated substitutions in BufA1 were grown in PYE medium supplemented or not with 225 μM $CuSO_4$.

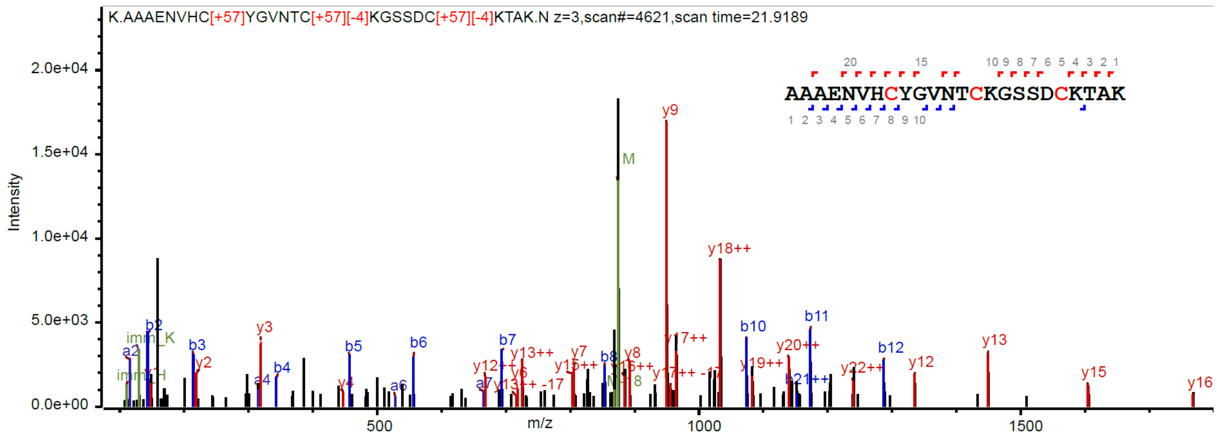


| Rt (s) | m/z | z | Modifications | E-value | Score | Delta [M+H] |
|---|---|---|---|---|---|---|
| 1315.25 | 874.0363 | 3 | 8C:57.02 14C:57.02 20C:57.02 | 4.2E-14 | 67.5 | -8.05825 |

| Best hit modified |
|---|
| AAAEVHCYGV[][N][T][C][-4,03]KGSS[D][C] ][-4,03]KT[A][K] |

**Figure S8. Bottom-up analysis of bufferin 2.** MS/MS spectra of the tryptic peptide containing the Cys$^{I}$, Cys$^{II}$ and Cys$^{III}$ residues, $^{5}$AAAENVHCYGVNTCKGSSDCKTA$^{28}$K ($[M+3H]^{3+}$ at *m/z* 874.04). Below is shown the hit proposed by the untargeted PTM search solution SpecGlobX.

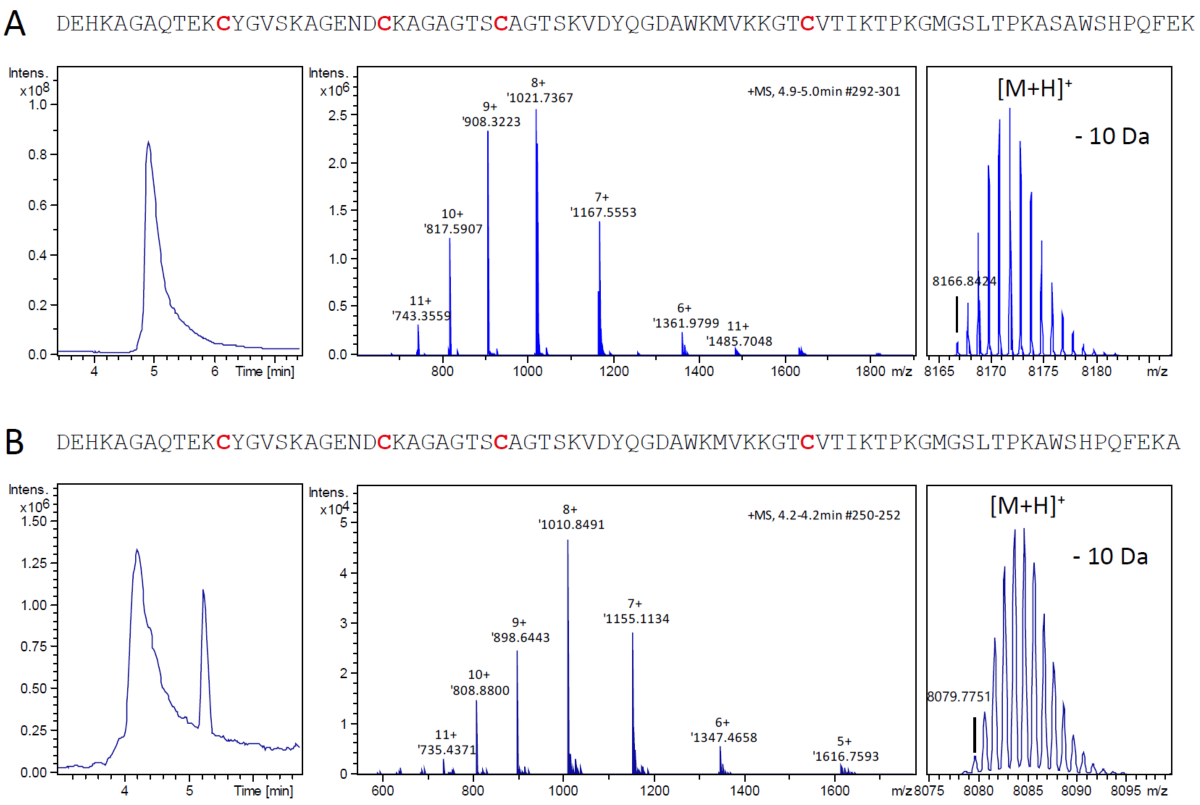


**Figure S9. Top-down analyses of purified bufferin 1.** Sequence of the peptides detected and LC-MS analyses of the cell extracts of *C. vibrioides* producing twin-strep-tagged bufferin 1 ($Buf1^{twstr}$) (A) or *E. coli* producing strep-tagged bufferin 1 ($Buf1^{str}$) (B). Total ion chromatograms, MS spectra and deconvoluted spectra are shown in the left, middle and right panels, respectively. For both peptides, -10 Da mass shifts were detected as compared to the unmodified sequences (monoisotopic Mw 8175.9108 Da for $Buf1^{twstr}$, Mw 8088.8788 Da for $Buf1^{str}$). Note that the twin-strep tag of $Buf1^{twstr}$ appears to undergo *in vivo* proteolytic cleavage in the middle of the tag, immediately after the first strep tag motif.

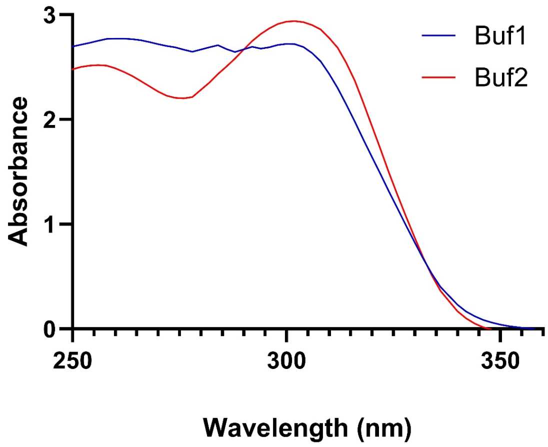


**Figure S10. Characterization of Buf1 and Buf2 by spectrophotometry**. Spectra of recombinant 6His-tagged Buf1 and Buf2. At similar protein concentrations $A^{305}$ is higher for $Buf2^{6\text{-}His}$ than for $Buf1^{6\text{-}His}$, consistent with Buf2 potentially harboring a larger number of heterocyclic PTMs.

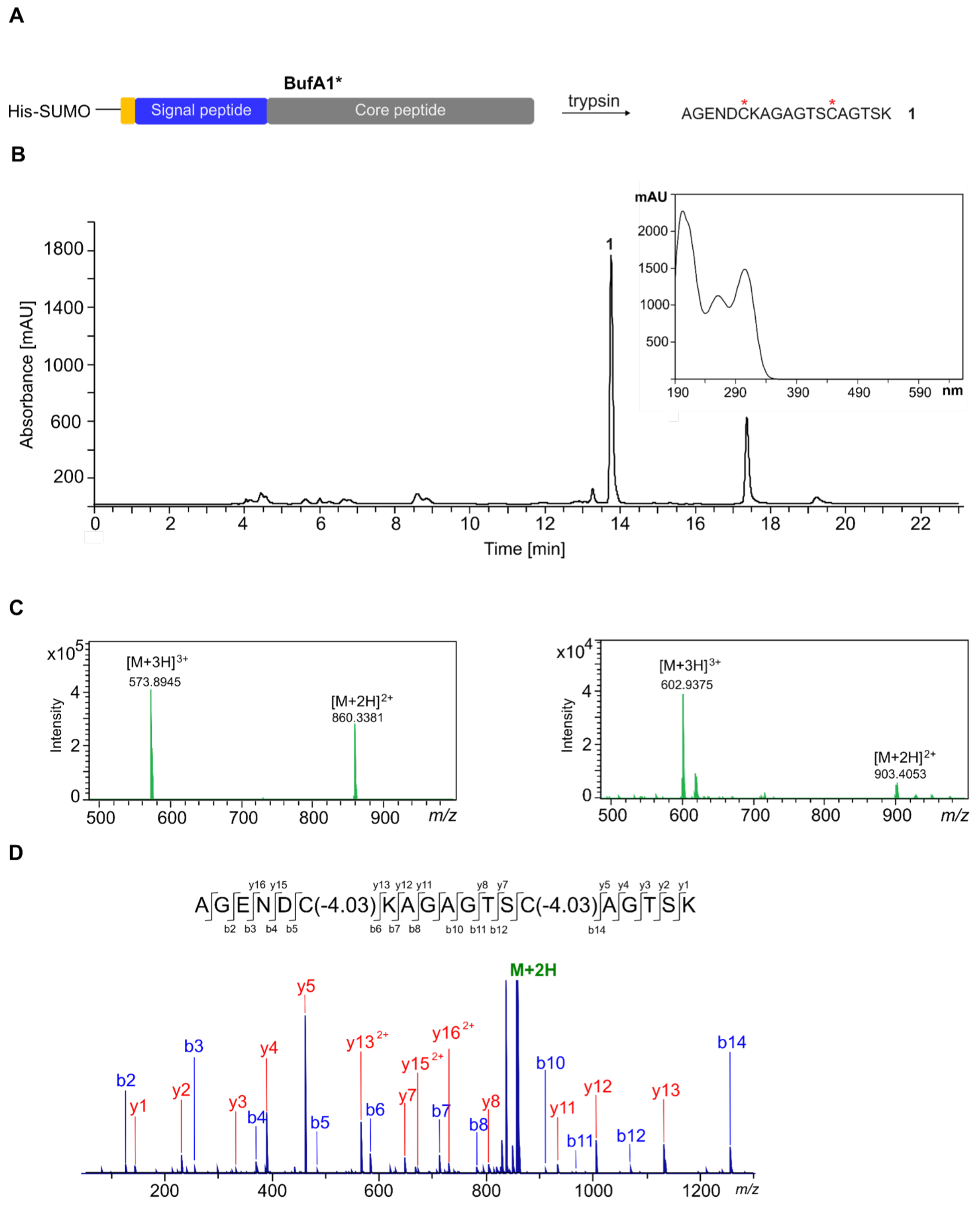


**Figure S11.** ***In vivo*** **production and purification of the central tryptic peptide.** A. Modified BufA1* was produced through co-expression of SUMO-BufA1 with BufB1 and BufC1 in *E. coli* and purified by Ni-affinity chromatography. The recombinant protein was digested with trypsin generating the 19-residue peptide 1 with modifications on the two Cys residues (marked with *). B. Chromatogram of HPLC purification (UV absorbance at 280 nm). The peak at RT = 13.85 min was collected and corresponds to peptide 1. The inset shows the corresponding UV/vis scan exhibiting a new absorbance maximum at 305 nm. C. Mass spectrum of purified unlabeled (left panel) and $^{13}C$, $^{15}N$-labeled peptide 1 (right panel). D. MS/MS spectrum of the central tryptic peptide ($[M+2H]^{2+}$ at *m*/z 860.3381).

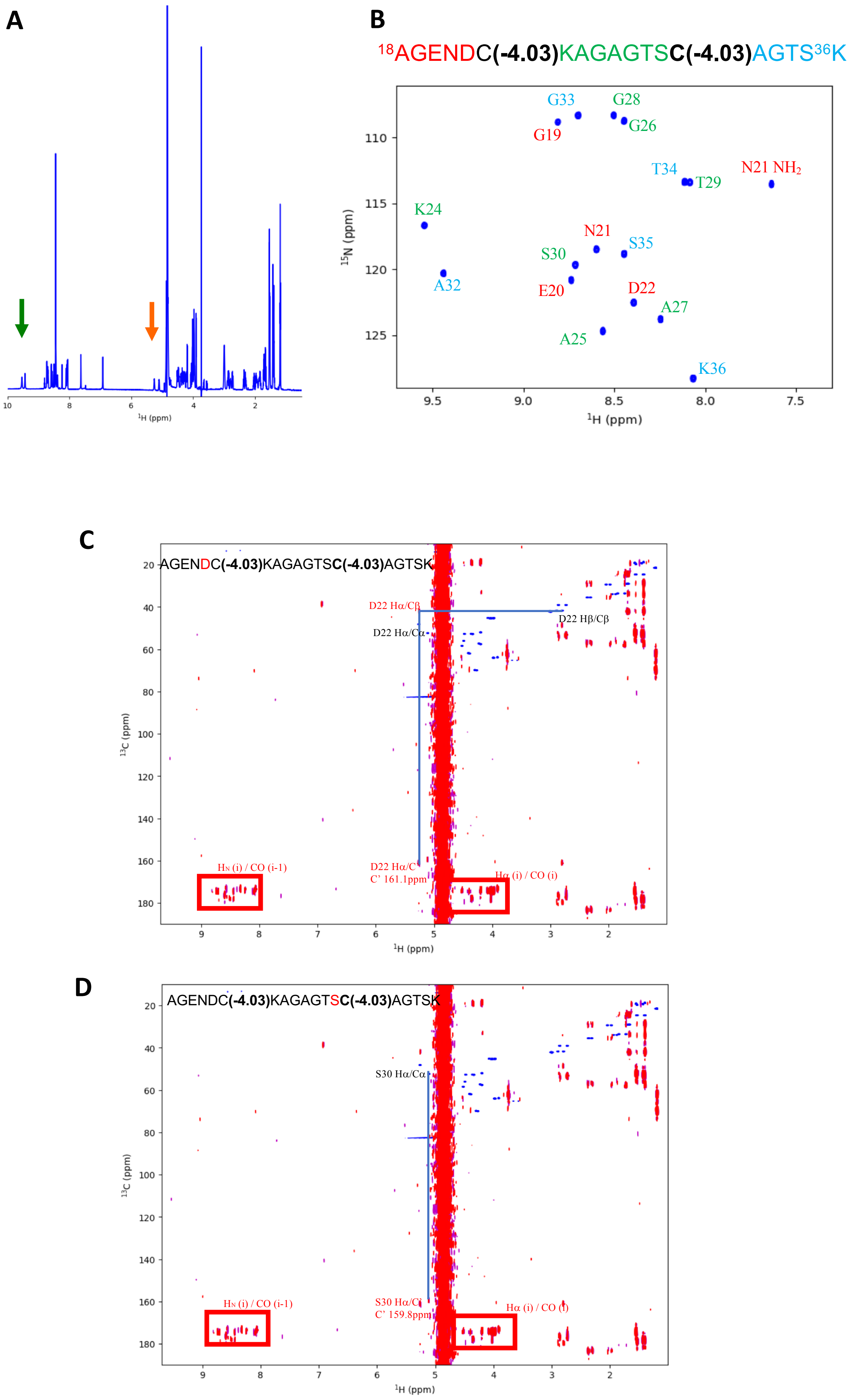


**Figure S12. NMR analyses of the central 19-mer tryptic peptide of Buf1. A**. $^{1}$H NMR spectrum of the peptide. Anomalous resonances are indicated by orange and green arrows. **B**. Natural abundance $^{1}$H, $^{15}$N HSQC spectrum. Assignments are reported in colored blocks according to the position of the amino acid residues with respect to the modified cysteine residues. **C** and **D**. $^{1}$H, $^{13}$C HSQC (blue) and HMBC (red) experiments on the 19-mer peptide at natural isotopic abundance.

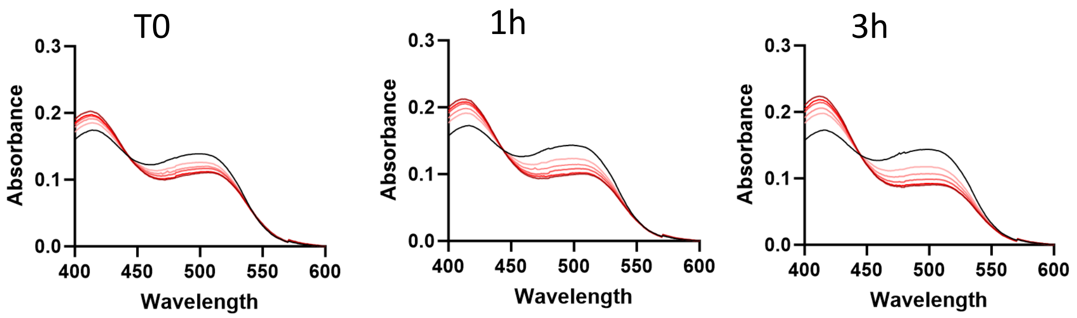


**Figure S13. Cu binding to bufferin 1 *in vitro*.** A competition spectroscopic $Cu^{2+}$ binding assay with Buf1 was performed as described (10). Increasing amounts of purified $Buf1^{twstr}$ (5 to 40 µM) were incubated with a colorimetric complex formed between $Cu^{2+}$ and 4-(2-pyridylazo)resorcinol (PAR) ($\varepsilon_{505nm}$ =41,500 $M^{-1}$ $cm^{-1}$), and the displacement of $Cu^{2+}$ from the $Cu^{2+}$-PAR complex by Buf1 was followed by spectrophotometry. The black curve represents the spectrum of the $Cu^{2+}$-PAR complex with no bufferin. The very slow formation of the Buf1-$Cu^{2+}$ complex suggests a partial displacement over time of an equilibrium between binding-incompetent and -competent forms of Buf1.

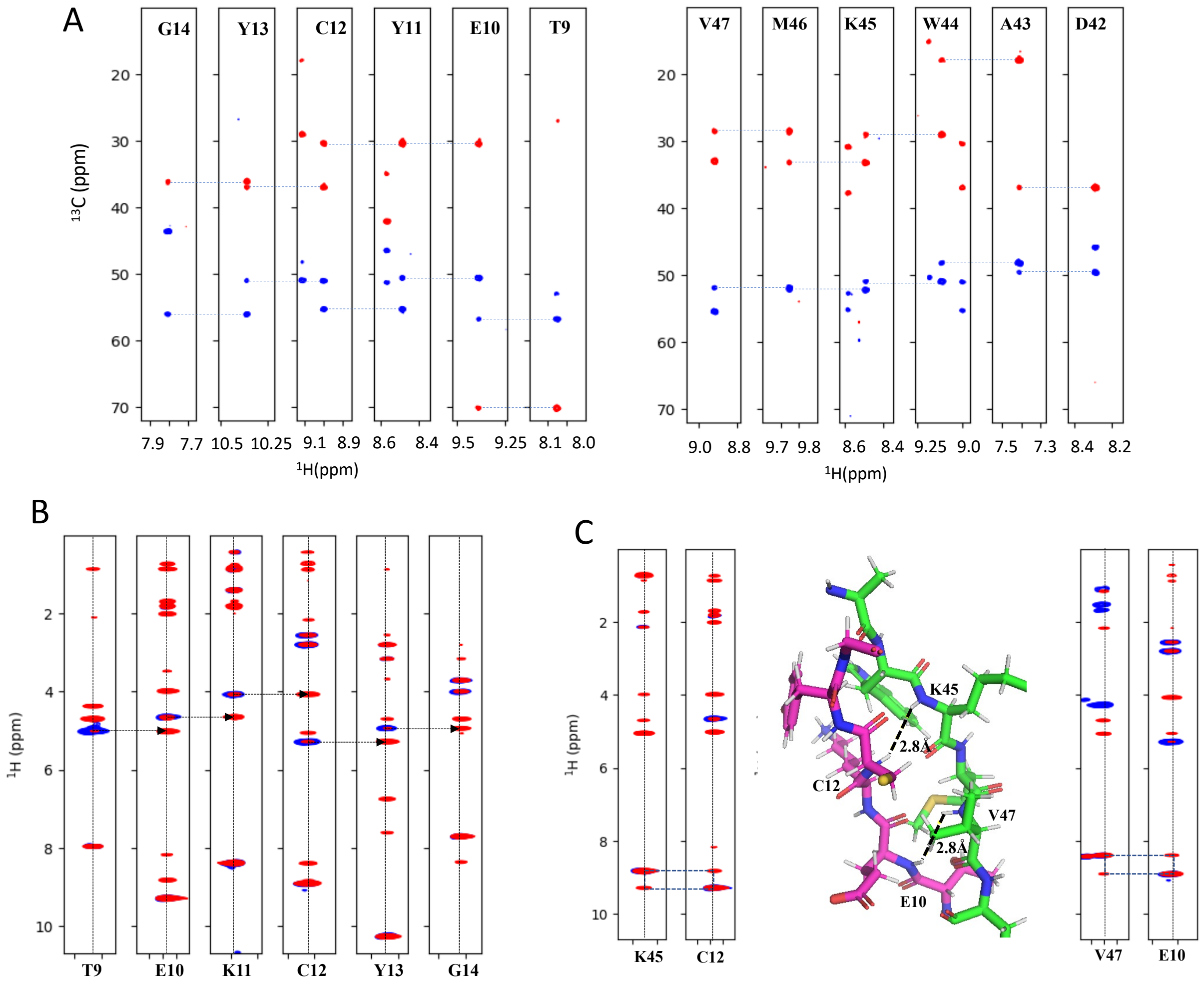


**Figure S14. Structural characterization of bufferin 1.** A. HSQC-NOESY and TOCSY experiments showing sequential walks through residues of the first and second β strands, respectively. B. Strips from TOCSY-HSQC (blue) and NOESY-HSQC (red) spectra of Buf1. Characteristic Hα(i-1) – HN(i) NOE contacts are indicated by arrows. These data confirm the assignement and demonstrate the extended nature of this segment. C. Strips through the TOCSY-HSQC (blue) and NOESY-HSQC (red) spectra of bufferin 1. These data show the inter-strand HN-HN NOE contacts, confirming the presence of an anti-parallel β sheet.

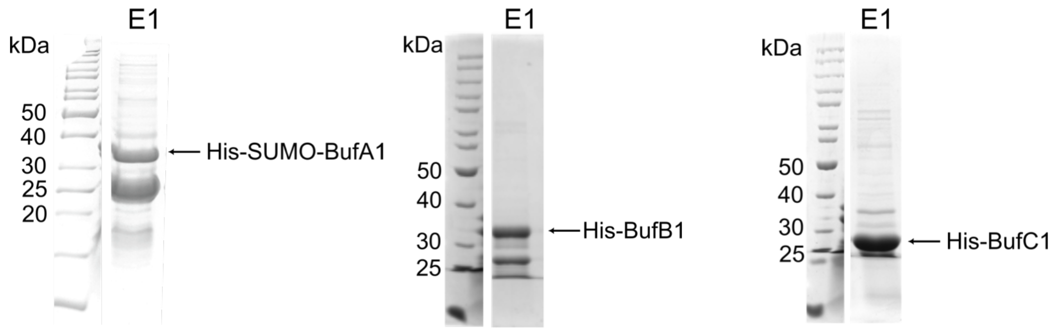


**Figure S15. Purification of the recombinant His-tagged proteins for the *in vitro* assay.** SDS-PAGE analyses of His-SUMO-BufA1 (left panel), His-BufB1 (middle panel) and His-BufC1 (right panel). E1 represents the elution fraction of each purification.

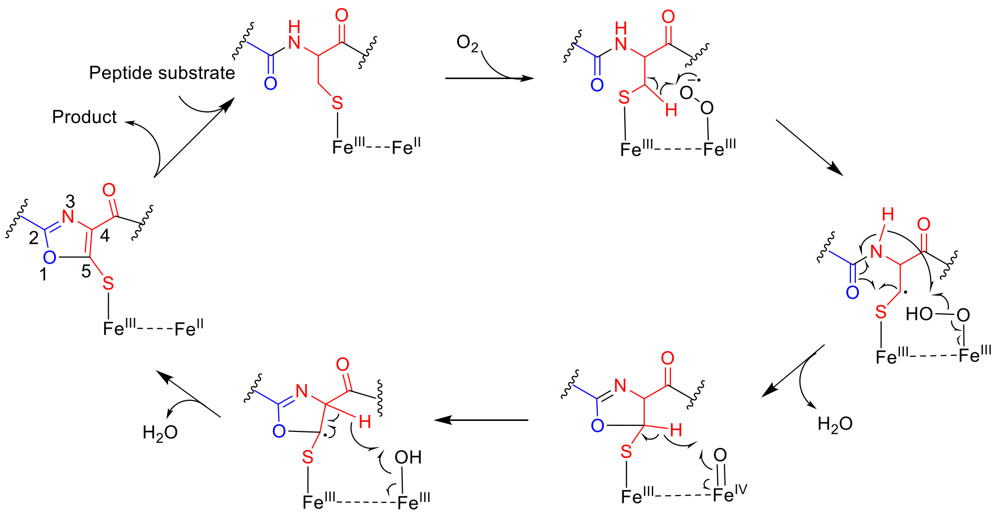


**Figure S16. Possible mechanism of BufB1-catalyzed conversion of Cys into a 5-thiooxazole motif involving a mixed valent di-iron center.** This process starts with $O_2$ activation and abstraction of a β-H followed by cyclization and ring oxidation.

### Supplementary Tables

***Table S1. Sequences of the precursor and core proteins used in this study***

BufA1_precursor
MTRTATTAALAAAIALSAGVSIAAADEHKAGAQTEKCYGVSKAGENDCKAGAGTSCAGTSKV
DYQGDAWKMVKKGTCVTIKTPKGMGSLTPKA

BufA1_core
DEHKAGAQTEKCYGVSKAGENDCKAGAGTSCAGTSKVDYQGDAWKMVKKGTCVTIKTPKGMG
SLTPKA

BufA2_precursor
MKETTMNSIKSVTLASAAALFALTSVAATPSFADSKKAAAENVHCYGVNTCKGSSDCKTAKN
ECKGQNECKGQGFKAMTKKACLTAGGSLTAPQ

BufA2_core
DSKKAAAENVHCYGVNTCKGSSDCKTAKNECKGQNECKGQGFKAMTKKACLTAGGSLTAPQ

BufA1_twin-strep_precursor (produced in *C. vibrioides*)
MTRTATTAALAAAIALSAGVSIAAADEHKAGAQTEKCYGVSKAGENDCKAGAGTSCAGTSKV
DYQGDAWKMVKKGTCVTIKTPKGMGSLTPKASAWSHPQFEKGGGSGGGSGGSAWSHPQFEK

BufA1_twin-strep_core
DEHKAGAQTEKCYGVSKAGENDCKAGAGTSCAGTSKVDYQGDAWKMVKKGTCVTIKTPKGMG
SLTPKASAWSHPQFEKGGGSGGGSGGSAWSHPQFEK

BufA1_twin-strep_core_truncated in vivo
DEHKAGAQTEKCYGVSKAGENDCKAGAGTSCAGTSKVDYQGDAWKMVKKGTCVTIKTPKGMG
SLTPKASAWSHPQFEK

BufA1_strep_precursor (produced in *E. coli*)
MTRTATTAALAAAIALSAGVSIAAADEHKAGAQTEKCYGVSKAGENDCKAGAGTSCAGTSKV
DYQGDAWKMVKKGTCVTIKTPKGMGSLTPKAWSHPQFEKA

BufA1_strep_core
DEHKAGAQTEKCYGVSKAGENDCKAGAGTSCAGTSKVDYQGDAWKMVKKGTCVTIKTPKGMG
SLTPKAWSHPQFEKA

BufA1_His-tag_TEV _core
DEHKAGAQTEKCYGVSKAGENDCKAGAGTSCAGTSKVDYQGDAWKMVKKGTCVTIKTPKGMG
SLTPKASSGENKYFQGGSHHHHHH

BufA2_His-tag_TEV_core
DSKKAAAENVHCYGVNTCKGSSDCKTAKNECKGQNECKGQGFKAMTKKACLTAGGSLTAPQS
SGNELYFQGGSHHHHHH

***Table S2. EPR parameters corresponding to the Easy-spin fit of the $Cu^{2+}$-Buf1 complex***

| g-factor | $A(^{63}Cu)$, [MHz] | $A(^{14}N)$, [MHz] | $A(^{14}N)$, [MHz] | $A(^{14}N)$, [MHz] | $A(^{14}N)$, [MHz] |
|---|---|---|---|---|---|
| 2.0400 | 23.60 | 55.68 | 51.06 | 64.48 | 64.36 |
| 2.0413 | 89.33 | 59.41 | 22.94 | 38.77 | 38.85 |
| 2.1856 | -604.72 | -9.44 | -24.89 | -4.28 | -4.27 |

***Table S3. EPR parameters corresponding to the Easy-spin fit of the $CuSO_4$ sample***

| g-factor | $A(^{63}Cu)$, [MHz] |
|---|---|
| 2.0874 | 9.33 |
| 2.0759 | 3.02 |
| 2.4159 | 389.89 |

***Table S4. Plasmids used in this study***

| Plasmids | Description | references |
|---|---|---|
| pCR4-TOPO | PCR cloning and sequencing | |
| pNPTS138 | *Neo sacB traJ* and *oriT*<br>Used for allelic replacement in *C vibrioides* | Gift of JY Matroule |
| pSRK-Km | Medium-copy number plasmid, Plac promoter | Gift of JY Matroule |
| pFUS2 | Promoterless *lacZ* for transcriptional fusions | (1) |
| pSigF | sigF sequence under the control of Plac in pSRK-Km | This work |
| pCA24-buf1A$^{str}$BCD | Expression of *buf1* in *E. coli* BL21 with strep-tagged A1 | This work |
| pCA24-psmCA | Expression of lasso peptide operon in *E. coli* BL21 | (5) |
| pETHisSUMO-A1 | Overexpression of BufA1 precursor fused to N-terminal His tag and SUMO in *E. coli* BL21(DE3) | This work |
| pACYC::HisB1 | Overexpression of BufB1 fused to N-terminal His tag in *E. coli* BL21(DE3) | This work |
| pACYC::HisC1 | Overexpression of BufC1 fused to N-terminal His tag in *E. coli* BL21(DE3) | This work |
| pACYC::HisC1B1Stag | Overexpression of both BufB1 with C-terminal Stag and BufC1with N-terminal His tag in *E. coli* BL21(DE3) | This work |

***Table S5. Recombinant strains used in this study***

| *Name* | *Genetic characteristics / use in this work* |
|---|---|
| NA1000 | Parental *C. vibrioides* strain |
| $Cv_{DKO}$ | Deletion of both *buf1* and *buf2* |
| $Cv_{buf1}$ | Harbours only *buf1* |
| $Cv_{buf2}$ | Harbours only *buf2* |
| $Cv_{buf1(A)}$ | Deletion of *bufB1-bufC1-doxX* genes in *buf1*: harbours only BufA1 precursor gene |
| $Cv_{buf2(A)}$ | Deletion of *bufB2-buf2* genes in *buf2*: harbours only BufA2 precursor gene |
| $Cv_{buf1(A\text{-}C\text{-}D)}$ | Deletion of *bufB1* gene in *buf2*: harbours *bufA1-bufC1-doxX* gene |
| $Cv_{buf1(A\text{-}B\text{-}D)}$ | Deletion of *bufC1* gene in *buf1*: harbours *bufA1-bufB1-doxX* genes |
| $Cv_{buf1(A\text{-}B\text{-}C)}$ | Deletion of *doxX* gene in *buf1*: harbours *bufA1-bufB1-bufC1* genes |
| $Cv_{buf1}^{TwStrep}$ | *buf1* with *bufA1* gene modified with a twin-strep tag-coding sequence at the 3’ end |
| $Cv_{buf1}^{CysIISer+CysIIISer\text{-}TwStrep}$ | *buf1* with *bufA1* gene carrying substitutions of $Cys^{II}$ and $Cys^{III}$ by Ser and a twin-strep tag-coding sequence at the 3’ end |
| $Cv_{buf1}^{CysIISer}$ | *buf1* with *bufA1* gene carrying substitutions of $Cys^{II}$ by Ser |
| $Cv_{buf1}^{CysIIISer}$ | *buf1* with *bufA1* gene carrying substitutions of $Cys^{III}$ by Ser |

| | |
|---|---|
| $Cv_{Bbuf1}^{CysIISer+CysIIISer}$ | *buf1* with *bufA1* gene carrying substitutions of $Cys^{II}$ and $Cys^{III}$ by Ser |
| $Cv_{buf1}^{CysISer+CysIVSer}$ | *buf1* with *bufA1* gene carrying substitutions of $Cys^{I}$ and $Cys^{IV}$ by Ser |
| $Cv_{buf1\text{-}lacZ}$ | Transcriptional fusions of *bufA1* with *lacZ* |
| $Cv_{buf2\text{-}lacZ}$ | Transcriptional fusions of *bufA2* with *lacZ* |
| *Ec* BL21(pCA24-buf1A$^{str}$BCD) | Expression of *buf1* (gain-of-function) in *E. coli* |
| *Ec* BL21(pCA24-psmCA) | Expression of control lasso peptide operon in *E. coli* |
| *Ec* BL21(*DE3,* pETHisSUMO-A1) | Production/purification of N-His$_6$-SUMO-BufA1 fusion |
| *Ec* BL21(*DE3*, pACYC::HisB1) | Production/purification of N-His$_6$-BufB1 for in vitro assay |
| *Ec* BL21(*DE3*, pACYC::HisC1) | Production/purification of N-His$_6$-BufC1 for in vitro assay |
| *Ec* BL21(*DE3*, pETHisSUMO-A1, pACYC::HisC1B1Stag) | Production/purification of N-His$_6$-SUMO-BufA1for trypsic digestion for NMR |

***Table S6. Pairs of PCR primers and synthetic genes.***

The restriction sites or the homologous regions for IVA or LIC cloning are underlined.

| | |
|---|---|
| Deletion of *buf1* (entire operon) | TAGAATTCGCGTCGATCAGATCATCCGTCC<br>ATTCTAGAGCGGGTCATGAGAGACTCTCC<br><br>TATCTAGACTGAGCCTGGACCATGTGACG<br>ATAAGCTTTATCGATTCCGCCCGGCAGGT |
| Deletion of *bufB1CD1* | ATGAATTCTCTTGCACCAGATCCTCCA<br>TATCTAGAAGGGGTCATGGTCTGATC<br><br>TATCTAGACTGAGCCTGGACCATGTGACG<br>ATAAGCTTTATCGATTCCGCCCGGCAGGT |
| Deletion of *buf2* (entire operon) | ATGCATGCGCCTTCTACGCCTTCGTGGTG<br>TATCTAGACTTGATGGAGTTCATGGTCGTT<br><br>TATCTAGAGAACCGCCAGCGTTGATCC<br>ATAAGCTTATGTTCCTTGTAGGCATAGGTGT |
| Deletion of *bufB2C2* | TAGCATGCATCCTGGCCTCGTGGAGC<br>TATCTAGAGAGAGTCATGGCCGCGTCCTC<br><br>TATCTAGAGAACCGCCAGCGTTGATCC<br>ATAAGCTTATGTTCCTTGTAGGCATAGGTGT |
| Deletion of *bufA1* | ATGAATTCGGCGTCGATCAGATCATCCGTC<br>TATCTAGAAGAGACTCTCCTGGTGGGTT<br><br>TATCTAGAGACCATCGTTCCCCGTCGCTT<br>ATAAGCTTACAGCGTCCAAACCACCTCC |
| Deletion of *bufB1* | ATGAATTCCTCTTGCACCAGATCCTCCACA<br>TATCTAGAAGGGGTCATGGTCTGATCTTCG<br><br>TATCTAGAGAGCCCGCGCATGTCTGA<br>ATAAGCTTGGGCTCACCGTGTCGAAC |
| Deletion of *bufC1* | ATGAATTCCTACATGGCGTGGGGCTGTCTCT<br>TATCTAGAGAAGGCCAGCAACTCAGACAT<br><br>TATCTAGAGACCTGGAAACGCAACCATG<br>ATAAGCTTGTCAGATCCATAAACCCGGTGT |
| Deletion of *bufD1* | ATGAATTCACGATACGCCTCAGGGTCTC<br>TATCTAGAGGTCATGGTTGCGTTTCCAG<br><br>TATCTAGACTGAGCCTGGACCATGTGACG<br>ATAAGCTTTATCGATTCCGCCCGGCAGGT |
| Deletion of *bufA2* | ATGAATTCCAGCTCGCGCCTCGTCA |

| | |
|---|---|
| | TATCTAGACAGCCATTACGCGACCGAC<br><br>ATTCTAGAGCGATAGCCTTTCCCGACGAC<br>ATGCATGCCAACAACTCACGCCCCTGACT ??? |
| Introduction of BufA1 variants in $Cv_{buf1(B-C-D)}$: flanking fragments (the central MluI-XbaI fragment is inserted between them) | TAGAATTCGCGTCGATCAGATCATCCGTCC<br>TAACGCGTTTGAAGGCGCTGATGCTACG<br><br>GCTCTAGAAGCCGTGCGAGCG<br>ATAAGCTTTATCGATTCCGCCCGGCAGGT |
| *bufA*1 fragment for reintroduction in $Cv_{buf1(B-C-D)}$ | CCGAATTCACGCGTCTCGGTGTCCGTCACGCC<br>GCTTCTAGAGCGGCGAGACGGG |
| Introduction of BufA1-twinstrep tagged: synthetic MluI-XbaI gene fragment | AAACGCGTCTCGGTGTCCGTCACGCCAGCCCCTAGTTCGCGTGCGAGGCC<br>ACGATGGTTACATCGTGTCTTCATCGCCCGAAACGGGCGGCCTGCGCGGC<br>GGTGAAATTTTGTTCGCCGATCACTGTAACCTCGGCCACCCCAGGAACGA<br>ACTCTCCCACGACCGAGCGGCGCAGGGCCGCCTCGGCGCGAACCCACCA<br>GGAGAGTCTCTCATGACCCGCACCGCCACCACCGCCGCCCTGGCCGCCGC<br>TATCGCTCTTTCGGCCGGCGTCTCCATCGCCGCCGCCGATGAGCACAAGG<br>CCGGCGCCCAGACCGAGAAGTGCTATGGCGTCTCCAAGGCCGGCGAGAA<br>CGACTGCAAGGCCGGCGCCGGCACCTCGTGCGCGGGCACTTCGAAGGTC<br>GACTACCAGGGCGACGCCTGGAAGATGGTCAAGAAGGGCACGTGCGTCA<br>CGATCAAGACGCCCAAGGGCATGGGCTCGCTGACGCCGAAGGCCTCGGC<br>CTGGTCGCACCCGCAGTTCGAGAAGGGCGGCGGCTCGGGCGGCGGCTCG<br>GGCGGCTCGGCCTGGTCGCACCCGCAGTTCGAGAAGTAACGCCGACCAT<br>CGTTCCCCGTCGCTTCCGGCCGCCCCCAAAGAGCGGTCGGGAGCGGCGC<br>GCCTCGAAGATCAGACCATGACCCCTTCCGCCGGCCTTGGCCTCAAATCG<br>CAGCACTATGGCGACGCGATCGCATGCGACGCCGAGGGCCTCTGGTTCG<br>AGGTTCATCCTGAAAACTACATGTCCGCCGGAGGGCCCCGTCTCGCCGCT<br>CTAGAAG |
| Mutagenesis of $Cys^{II}$ in BufA1 | CGGCGAGAACGACAGCAAGGCCGGCGC<br>GCGCCGGCCTTGCTGTCGTTCTCGCCG |
| Mutagenesis of $Cys^{III}$ in BufA1 | CGCCGGCACCTCGT CGGCGGGCACTTCGAA<br>TTCGAAGTGCCCGC CGACGAGGTGCCGGCG |
| Introduction of $buf1^{CysISer+CysIVSer}$: synthetic MluI-XbaI gene fragment | AAACGCGTCTCGGTGTCCGTCACGCCAGCCCCTAGTTCGCGTGCGAGGCC<br>ACGATGGTTACATCGTGTCTTCATCGCCCGAAACGGGCGGCCTGCGCGGC<br>GGTGAAATTTTGTTCGCCGATCACTGTAACCTCGGCCACCCCAGGAACGA<br>ACTCTCCCACGACCGAGCGGCGCAGGGCCGCCTCGGCGCGAACCCACCA<br>GGAGAGTCTCTCATGACCCGCACCGCCACCACCGCCGCCCTGGCCGCCGC<br>TATCGCTCTTTCGGCCGGCGTCTCCATCGCCGCCGCCGATGAGCACAAGG<br>CCGGCGCCCAGACCGAGAAGAGCTATGGCGTCTCCAAGGCCGGCGAGAA<br>CGACTGCAAGGCCGGCGCCGGCACCTCGTGCGCGGGCACTTCGAAGGTC<br>GACTACCAGGGCGACGCCTGGAAGATGGTCAAGAAGGGCACGAGCGTCA<br>CGATCAAGACGCCCAAGGGCATGGGCTCGCTGACGCCGAAGGCCTAACG<br>CCGACCATCGTTCCCCGTCGCTTCCGGCCGCCCCCAAAGAGCGGTCGGGA<br>GCGGCGCGCCTCGAAGATCAGACCATGACCCCTTCCGCCGGCCTTGGCCT<br>CAAATCGCAGCACTATGGCGACGCGATCGCATGCGACGCCGAGGGCCTC<br>TGGTTCGAGGTTCATCCTGAAAACTACATGTCCGCCGGAGGGCCCCGTCT<br>CGCCGCTCTAGAAG |
| Insertion of BufA2 variant in<br>variants in $Cv_{buf2(B-C)}$ : flanking fragments (the central SacI-XbaI fragment is inserted between them) | ATGAATTCCAGCTCGCGCCTCGTCA<br>TGAGCTCCGGTGACCAGGGTGAC<br><br>TGAGCTCCGGTGACCAGGGTGAC<br>ATGCATGCCAACAACTCACGCCCCTGACT ??? |
| Introduction of BufA1-$His_6$ tagged: synthetic MluI-XbaI gene fragment | AAACGCGTCTCGGTGTCCGTCACGCCAGCCCCTAGTTCGCGTGCGAGGCC<br>ACGATGGTTACATCGTGTCTTCATCGCCCGAAACGGGCGGCCTGCGCGGC<br>GGTGAAATTTTGTTCGCCGATCACTGTAACCTCGGCCACCCCAGGAACGA<br>ACTCTCCCACGACCGAGCGGCGCAGGGCCGCCTCGGCGCGAACCCACCA<br>GGAGAGTCTCTCATGACCCGCACCGCCACCACCGCCGCCCTGGCCGCCGC<br>TATCGCTCTTTCGGCCGGCGTCTCCATCGCCGCCGCCGATGAGCACAAGG<br>CCGGCGCCCAGACCGAGAAGTGCTATGGCGTCTCCAAGGCCGGCGAGAA |

| | |
|---|---|
| | CGACTGCAAGGCCGGCGCCGGCACCTCGTGCGCGGGCACTTCGAAGGTC<br>GACTACCAGGGCGACGCCTGGAAGATGGTCAAGAAGGGCACGTGCGTCA<br>CGATCAAGACGCCCAAGGGCATGGGCTCGCTGACGCCGAAGGCCAGCTC<br>GGGCGAGAACCTGTACTTCCAGGGCGGCTCGCATCACCATCACCATCACT<br>AACGCCGACCATCGTTCCCCGTCGCTTCCGGCCGCCCCCAAAGAGCGGTC<br>GGGAGCGGCGCGCCTCGAAGATCAGACCATGACCCCTTCCGCCGGCCTT<br>GGCCTCAAATCGCAGCACTATGGCGACGCGATCGCATGCGACGCCGAGG<br>GCCTCTGGTTCGAGGTTCATCCTGAAAACTACATGTCCGCCGGAGGGCCC<br>CGTCTCGCCGCTCTAGAAG |
| Introduction of BufA2-$His_6$ tagged: synthetic SacI-XbaI gene fragment | CGGAGCTCAGTACTGGGAAGCGGCCCGCAAGGCGCTGGTCTAGGCGGGG<br>CGGCTGTCGCCAGGAGGGGCTCCCCGGTACCCTCTTCACCGAGCACAGC<br>AAAAGGGTCCGGTCCTCCGCCAGATCACCAGGATCCGGTGTCCCGCGAC<br>TTTGTCCGAGTTTTTTGAGTATGGTCTGTAACCTTGCCGGGGTCGGTCGCG<br>TAATGGCTGATGAGGCGGGCCGATGGACGGCTTGGACCTCCTGTGAAGG<br>AAACGACCATGAACTCCATCAAGAGCGTCACCCTGGCCTCGGCCGCCGC<br>CCTGTTCGCCCTGACCAGCGTCGCCGCTACCCCCAGCTTCGCCGACAGCA<br>AGAAGGCCGCCGCTGAAAACGTTCACTGCTACGGCGTGAACACCTGTAA<br>GGGCAGCTCGGACTGCAAGACCGCCAAGAACGAGTGCAAGGGTCAGAAT<br>GAGTGCAAGGGTCAGGGCTTCAAGGCCATGACCAAGAAGGCTTGCCTCA<br>CCGCCGGCGGTTCGCTGACCGCTCCCCAGTCGAGCGGCGAGAACCTGTA<br>CTTCCAGGGCGGCTCGCATCACCATCACCATCACTAAGGCGTGCGGACGC<br>GACGACGCAGACGTCTGGCGATAGCCTTTCCCGACGACACGTCGTCGCGT<br>CTTATCCTCCCATGGGCCAATCAAGTGAGGACGCGGCCATGACTCTCCAG<br>CCATTCGACGGCTTCGGTCTGGGCCTGCGCCCGCCGCACTATCGCGCGTT<br>TCTAGACA |
| Transcriptional fusion with *buf1A* | TAAAGCTTCCAGGAAGTCGACATGGTCCTC<br>TAGGATCCGCACGTGCCCTTCTTGACCATC |
| Transcriptional fusion with *buf2A* | TAAAGCTTCCTGATCCTGTTCCGGGAGT<br>TAGCATGCAGCCTTCTTGGTCATGGCCTT |
| Cloning of *sigF* | TACATATGACGGACACCGAGACCCGA<br>ATAAGCTTGGTCCCCCGCCCTGCATCG |
| Cloning of *buf1* in pCA24 | TTATAAGATCTTTCATTAAAGAGGAGAGTCTCTCATGACCCGCAC<br>TTATAAAGCTTCAGGGCGTTCGCGTCACATGG |
| *buf1* with BufA1-Strep tag in pCA24 | TTATAAGATCTTTCATTAAAGAGGAGAGTCTCTCATGACCCGCAC<br>TTTAGGCTTTTTCAAACTGCGGATGGCTCCAGGCCTTCGGCGTCAGCGAG |
| IVA plasmid backbone of pACYC Duet-1 (MCS1) | GCGGCCGCATAATGCTTAAG<br>CGAATTCGGATCCTGGCTGTG |
| IVA for *bufB1* in MCS1 of pACYCDuet-1 | CAGGATCCGAATTCGGATCAGACCATGACCCCTTCCG<br>GCATTATGCGGCCGCGCCAGCAACTCAGACATGCG |
| IVA for *bufC1* in MCS1 of pACYCDuet-1 | CAGGATCCGAATTCGATGTCTGAGTTGCTGGCCTTC<br>GCATTATGCGGCCGCTCATGGTTGCGTTTCCAGGTC |
| IVA plasmid backbone of pACYC Duet-1 (MCS2 with S-tag) | GGTACCCTCGAGTCTGGTAAAG<br>CATATGTATATCTCCTTCTTATAC |
| IVA for *bufB1* in MCS2 of pACYCDuet-1 | GGA GAT ATA CAT ATG ATG ACC CCT TCC GCC GGC<br>AGA CTC GAG GGT ACC GAC ATG CGC GGG CTC GCG |
| LIC for *bufA1* in pETSUMO | TACTTCCAATCCAATGCAATGACCCGCACCGCCACCAC<br>TTATCCACTTCCAATGTTATTATTAGGCCTTCGGCGTCAGCG |
